# Structural basis of TMPRSS11D specificity and autocleavage activation

**DOI:** 10.1101/2024.10.09.617371

**Authors:** Bryan J. Fraser, Ryan P. Wilson, Olzhas Ilyassov, Jackie Lac, Aiping Dong, Yen-Yen Li, Alma Seitova, Yanjun Li, Zahra Hejazi, Tristan M.G. Kenney, Linda Z. Penn, Aled Edwards, Gregg B. Morin, François Bénard, Cheryl H. Arrowsmith

**Affiliations:** Structural Genomics Consortium Toronto, Toronto, Ontario Canada; Department of Medical Biophysics, University of Toronto, Toronto, Ontario Canada Department of Radiology, University of British Columbia, Vancouver, British Columbia, Canada; Princess Margaret Cancer Centre, Toronto, Ontario Canada; Canada’s Michael Smith Genome Sciences Centre, Vancouver, British Columbia Canada; British Columbia Cancer Research Institute, Vancouver, British Columbia Canada; University of British Columbia, Vancouver, British Columbia Canada

## Abstract

Transmembrane Protease, Serine-2 (TMPRSS2) and TMPRSS11D are human proteases that enable SARS-CoV-2 and Influenza A/B virus infections, but their biochemical mechanisms for facilitating viral cell entry remain unclear. We demonstrate these proteases can spontaneously and efficiently cleave their own zymogen activation motifs, thereby activating their wider protease activity on other cellular substrates. We determined TMPRSS11D co-crystal structures in complexes with a native TMPRSS11D zymogen activation motif and with an engineered activation motif, providing insights into TMPRSS11D autocleavage activation and revealing unique regions of its substrate binding cleft. We further show that a protease inhibitor that underwent clinical trials for TMPRSS2-targeted COVID-19 therapy, nafamostat mesylate, was rapidly cleaved by TMPRSS11D and converted to low activity derivatives. These insights into human protease viral tropism and into liabilities with existing human serine protease inhibition strategies will guide future drug discovery campaigns for these targets.

## INTRODUCTION

Human respiratory viruses pose significant threats to global public health. The emergence of Severe Acute Respiratory Syndrome Coronavirus-2 (SARS-CoV-2) and the COVID-19 pandemic has highlighted the urgent need for antiviral therapeutics that can be deployed when respiratory virus vaccines are not available. To this end, the viral entry mechanism of SARS-CoV-2 has been intensely studied to identify druggable protein targets for SARS-CoV-2 and other human coronavirus infections^1–5^. A family of cell surface human proteases, the Type II Transmembrane Serine Proteases (TTSPs), have been shown to drive efficient SARS-CoV-2 viral entry and infection and are important drug targets for host-targeted antiviral prophylactics and/or therapeutics^3,5–8^.

TTSPs are first produced as inactive precursors (zymogens). Proteolytic cleavage at a specific (Arg/Lys)-(Ile/Val) peptide bond in their activation motif activates their serine protease (SP) domains, enabling them to proteolyze cellular substrates^9–14^ (Fig. 1a). Thus, zymogen cleavage activation is the most significant post-translational modification for TTSP enzymatic activity and biological function. Functionally important substrates of the SP domains of TTSPs include membrane proteins, extracellular matrix proteins, hormone precursors, protease zymogens and viral particles (Fig. 1b)^15^. One of the most intensely studied TTSPs, Transmembrane protease, serine-2 (TMPRSS2), cleaves the SARS-CoV-2 Spike protein to enable viral infection^3,8,16–18^. Other TTSPs have also been shown to play critical roles in SARS-CoV-2 and influenza virus infections in the absence of TMPRSS2^19–21^. TMPRSS11D (Human Airway Trypsin-like protease; HAT; Uniprot O60235), a member of the HAT/DESC subfamily of TTSPs, is highly expressed in the human airways and can enable SARS-CoV-2 and Influenza A infection^19,22–24^. The protein domain organization of TMPRSS11D (and all other HAT/DESC subfamily members) consists of a small cytoplasmic domain at the N-terminus, a single-pass transmembrane domain, a Sea urchin, Enteropeptidase and Agrin (SEA) domain, and a C-terminal SP domain. The only reported protein crystal structure of the HAT/DESC subfamily is the SP domain of TMPRSS11E (DESC1; Uniprot Q9UL52)^25^ and no selective inhibitors of any HAT/DESC subfamily members have been described to date. Furthermore, the high sequence similarity shared amongst the HAT/DESC group suggests that they may have similar protein substrate preferences, and it is unclear what interconnected role(s) these proteases have and any potential redundancy they have within the human airways.

**Figure 1.**
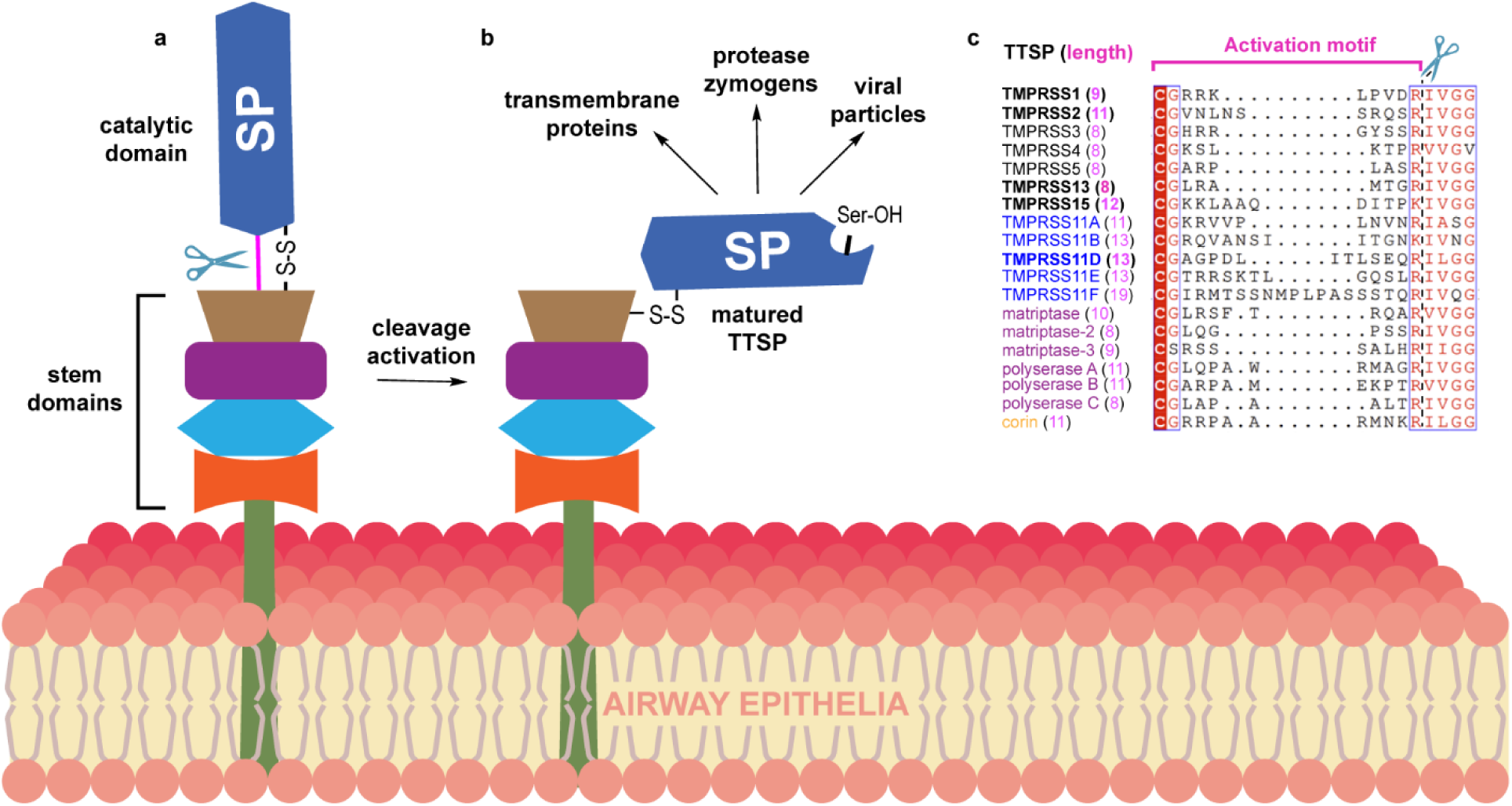
The zymogen activation motif of TTSPs are cleaved by trypsin-like serine proteases and their residue composition is unique to each TTSP. **a**, Schematic of an inactive (zymogen) TTSP at the cell surface. The catalytic Serine Protease (SP) domain is connected to the non-catalytic (stem) domains through a disulfide bond (S-S) and the zymogen activation motif peptide bond, shown as a pink line. The zymogen motif peptide bond is cleaved (indicated with scissors) to form (**b**) the matured TTSP that has enzymatic activity and can cleave protein and/or peptide substrates. c, Multiple sequence alignment of the zymogen activation motif of all human TTSPs. TTSPs are colored by TTSP subfamily; hepsin/TMPRSS-black; HAT/DESC-blue; matriptase-magenta; corin-orange. The length of the zymogen activation motif is indicated in parentheses for each TTSP. Scissors and a dashed black line indicate where TTSPs are cleaved during protease zymogen activation.

In this study, we applied a protein engineering method that enabled the high-yield production of active TMPRSS11D protease and determined the first X-ray crystal structures of the TMPRSS11D serine protease domain bound to a native, cleaved product molecule. We investigated the zymogen activation motifs of TMPRSS2 and TMPRSS11D, and demonstrated that these proteases recognize their own zymogen activation motifs to turn on their proteolytic activity, potentially explaining why their protease activities are exploited by respiratory viruses for viral entry. Through a combination of biochemical and biophysical assays, crystal structures, and computational modeling, we gained insights into TMPRSS11D substrate recognition and nafamostat inhibitor binding kinetics. Furthermore, the structural characterization of TMPRSS11D complexed with peptide ligands lays the groundwork for the rational design of targeted inhibitors aimed at modulating its activity and attenuating respiratory viral infections, including those caused by emerging pathogens such as SARS-CoV-2.

## RESULTS

### TMPRSS2 and TMPRSS11D cleave their own zymogen activation motifs, hindering recombinant protein overexpression

We previously acquired a highly active source of the TMPRSS2 ectodomain using a directed activation strategy (das) that replaced the TMPRSS2 zymogen activation motif with DDDDK^255^↓IVGG (dasTMPRSS2). This strategy enables control of cleavage activation after protein purification from baculovirus-infected Sf9 insect cells^10^. Without the DDDDK sequence replacement or a catalytic serine mutation (S441A), we were unable to overexpress any soluble TMPRSS2 protein. Others have since replicated production and purification of the dasTMPRSS2 protein to determine various TMPRSS2 crystal structures^26,27^. We reapplied this approach to TMPRSS11D by replacing its zymogen activation motif, LSEQR^186^↓ILGG with DDDDK^186^↓ILGG to create a dasTMPRSS11D construct, which greatly improved protein expression levels relative to the wild-type TMPRSS11D ectodomain protein which showed no detectable protein band (eTMPRSS11D; Fig. 2a). To study the TMPRSS11D ectodomain with its native zymogen activation motif, we introduced a catalytic serine mutation, S368A, which improved protein expression levels (eTMPRSS11D S368A; Fig. 2b). These protein expression data indicated that constitutive TMPRSS11D protease activity poses challenges to its overexpression in recombinant host cells, similar to TMPRSS2^10^.

**Figure 2.**
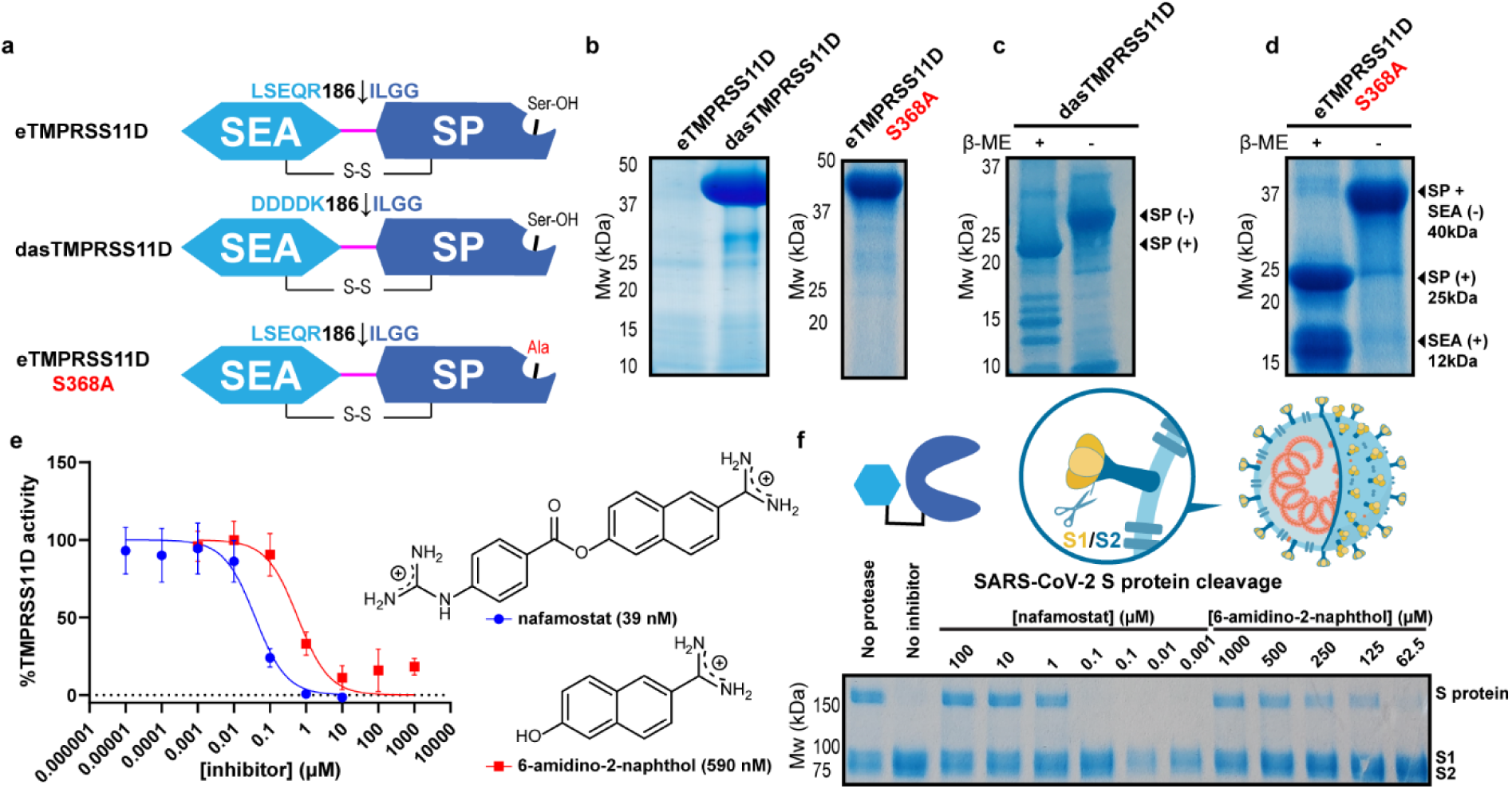
Soluble and proteolytically active TMPRSS11D is accessible by replacing its zymogen activation motif with a DDDDK sequence. **a**, Schematics of soluble human TMPRSS11D protein constructs that span the TMPRSS11D ectodomain. The two protein domains include the Sea urchin, Enteropeptidase and Agrin (SEA) and Serine Protease (SP) domains which are connected by a disulfide bond (S-S) and the R186-I187 peptide bond. Mutations targeting the R186-I187 cleavage site for each protein construct are indicated. **b**, TMPRSS11D protein test expression studies from baculovirus-infected Sf9 insect cells. The indicated TMPRSS11D protein was purified from media through IMAC purification and protein content evaluated by SDS-PAGE. Samples were thermally denatured and reduced (4x Laemmeli buffer containing 5 mM β-mercaptoethanol, 95°C, 5 min) prior to gel separation. **c**, Purified, active dasTMPRSS11D protein. SDS-PAGE samples were thermally denatured and reduced (+) or were not heated and not reduced (−) in advance of gel separation. **d**, Purified, activated eTMPRSS11D S368A protein. **e**, TMPRSS11D half-maximal inhibitory concentration (IC_50_) plots for the indicated small molecule inhibitors. Peptidase assays contained 0.2 nM dasTMPRSS11D and 100 µM Boc-QAR-AMC substrate and relative protease activity determined across the first 60 seconds of the reaction after substrate addition. Inhibitors were pre-incubated with dasTMPRSS11D for 15 minutes prior to the start of the assay. **f**, SARS-CoV-2 S protein cleavage assay with dasTMPRSS11D protease and S protein substrate. Purified recombinant S protein was incubated with 10 nM dasTMPRSS11D protease for 15 minutes at 20°C. Samples were prepared for SDS-PAGE through addition of 4X Laemelli buffer and heated (95°C for 5 minutes), then separated on a gel and stained with Coomassie blue.

When the dasTMPRSS11D protein was overexpressed, purified, and concentrated to ∼5 mg/mL, it underwent rapid autocleavage activation (Fig. 2c). The activated dasTMPRSS11D protein migrated on SDS-PAGE gels at a molecular weight of approximately 27 kDa and produced a single elution peak when purified further by size-exclusion chromatography (Extended Data Fig. 1a). These data suggested that the dasTMPRSS11D protein had released its SEA domain through proteolytic cleavage. In contrast, the eTMPRSS11D S368A protein did not undergo autocleavage activation at a protein concentration of 5 mg/mL. However, overnight incubation of eTMPRSS11D S368A with nanomolar amounts of active dasTMPRSS11D produced eTMPRSS11D S368A protein bands migrating at 25 kDa and 12 kDa when SDS-PAGE samples were reduced, and a single protein band at 40 kDa when the sample was not reduced prior to gel separation (Fig. 2d). These banding patterns suggest that the cleaved eTMPRSS11D S368A protein contained its SEA and SP domains, and the dasTMPRSS11D protease successfully activated the eTMPRSS11D S368A protein sample at its LSEQR^186^↓ILGG zymogen activation motif. We repeated this experiment for TMPRSS2 and found that eTMPRSS2 S441A can be cleaved at its SRQSR^255^↓IVGG zymogen activation motif through incubation with trace amounts of dasTMPRSS2 (Extended Data Fig. 1b-c). Thus, these overexpression studies and protease activation assays provide direct biochemical evidence that TMPRSS2 and TMPRSS11D are capable of autocleavage activation, and that protease autocleavage activation can pose challenges in their recombinant overexpression.

### dasTMPRSS11D protease activity is disabled by small molecule inhibitors

TMPRSS11D has been identified as a potential antiviral drug target, motivating us to establish inhibitor screening assays and detailed kinetic experiments. The activated dasTMPRSS11D protein displayed robust peptidase activity towards the fluorogenic peptide substrates Boc-QAR-AMC and Boc-QRR-AMC with respective *K*_*m*_ values of 8.3 µM and 181 µM (Extended Data Fig. 2). The Boc-QAR-AMC and Boc-QRR-AMC substrates exhibited signs of weak substrate inhibition, with apparent inhibition constants (*K_i,app_s*) of 1.4 mM and 2.1 mM, respectively. The Boc-QAR-AMC substrate was selected to measure dasTMPRSS11D inhibition by small molecule protease inhibitors. Nafamostat and 6-amidino-2-naphthol inactivated or inhibited dasTMPRSS11D with half-maximal inhibitory concentrations (IC_50_s) of 39 nM and 590 nM, respectively, when the compounds were preincubated with dasTMPRSS11D for 15 minutes prior to initiating the assay (Fig. 2e).

The dasTMPRSS11D protease cleaved SARS-CoV-2 S protein as a substrate, converting the 150 kDa molecular weight protein band into proteins migrating at 100 kDa and 70 kDa, respectively (Fig. 2f). When dasTMPRSS11D was preincubated with nafamostat or 6-amidino-2-naphthol, the protease activity was blocked as indicated by the presence of the intact SARS-CoV-2 S protein substrate protein. These trends matched the potencies obtained in Boc-QAR-AMC peptidase assays and confirmed that this orthogonal SARS-CoV-2 S protein cleavage assay could be used for hit compound confirmation after high-throughput TMPRSS11D drug screening campaigns.

### Nafamostat acylates dasTMPRSS11D, but rapidly hydrolyzes

Nafamostat and other ester-based serine protease inhibitors rapidly acylate the conserved catalytic serine residue to block enzymatic activity, then the acyl-enzyme complex eventually hydrolyzes to restore proteolytic activity (Fig. 3a). After hydrolysis of the acyl-enzyme complex, the product molecules have low inhibitory potency towards the protease relative to the starting ester compound^28^. We hypothesized that the same nafamostat inhibition mechanism applies to TMPRSS11D. To determine these parameters, nafamostat and Boc-QAR-AMC substrate were added simultaneously to wells containing dasTMPRSS11D (Fig. 3b). The reaction progress curves plateaued over time, indicating nafamostat acylates TMPRSS11D’s S368 residue. We used progress curve fitting to determine a TMPRSS11D *k*_inact_/K_I_ value of 0.094 µM^−1^min^−1^ for nafamostat. Interestingly, the nafamostat:TMPRSS11D acyl-enzyme complex rapidly hydrolyzed and restored dasTMPRSS11D peptidase activity, with an inhibition half-life (*t*_1/2_) of only 0.2 hours (Fig. 3c). In comparison, the TMPRSS2 *k*_inact_/K_I_ and inhibition *t*_1/2_values for nafamostat were previously determined to be 180 µM^−1^min^−1^ and 14.7 hours, respectively^10^. Thus, nafamostat more potently inactivates TMPRSS2 protease activity than TMPRSS11D and the TMPRSS2 inhibition is retained over 73-fold longer than for TMPRSS11D.

**Figure 3.**
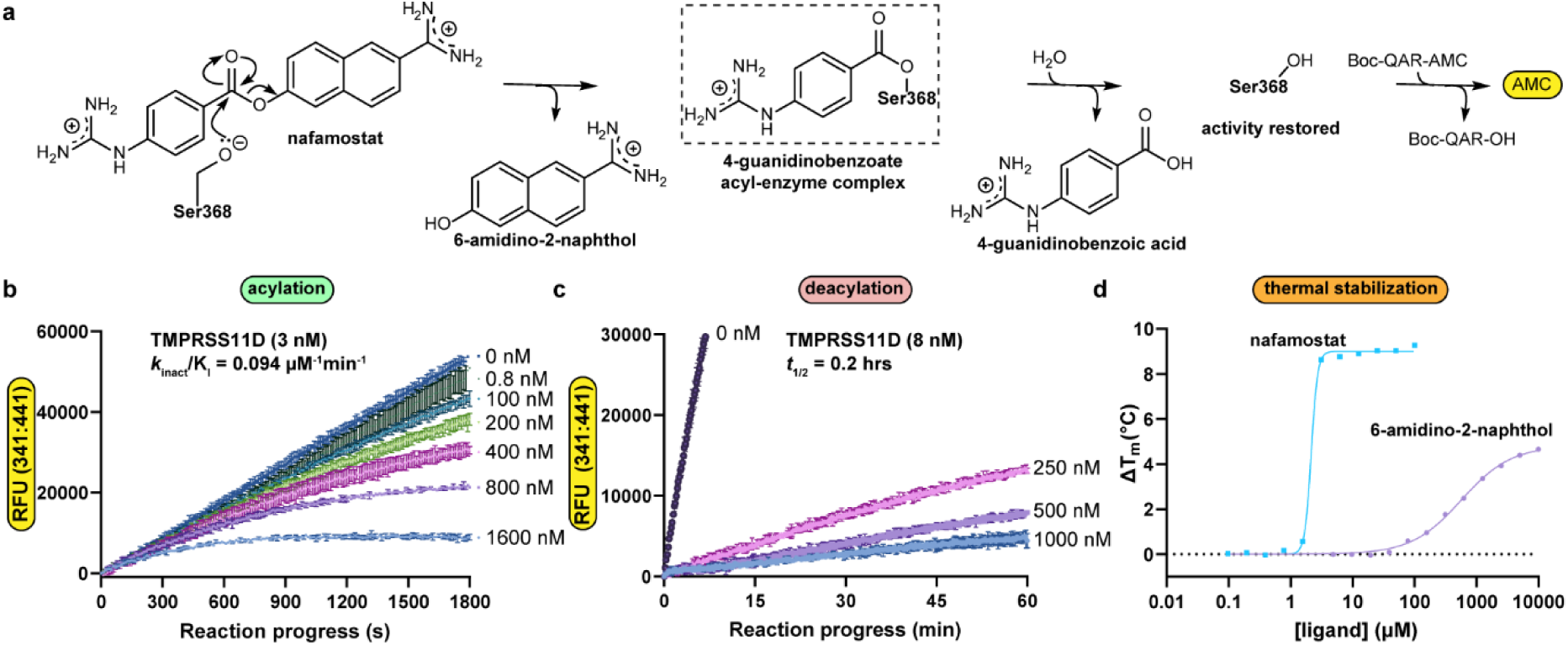
Nafamostat rapidly acylates dasTMPRSS11D, then hydrolyzes to restore protease activity. **a**, The putative nafamostat covalent inhibition mechanism for TMPRSS11D. **b**, Peptidase activity progress curves of dasTMPRSS11D (3 nM) with nafamostat at the indicated inhibitor concentrations added simultaneously with Boc-QAR-AMC substrate (100 µM final). **c**, Peptidase activity progress curves of dasTMPRSS11D (8 nM) pre-incubated (10 min) with the indicated concentrations of nafamostat before being transferred to wells containing Boc-QAR-AMC substrate (100 µM final). **d**, Melting temperature shifts (ΔT_m_s) of dasTMPRSS11D protein in the presence of the indicated concentrations of nafamostat (teal datapoints) or 6-amidino-2-naphthol (violet datapoints) ligands. Each assay contained 2 µg dasTMPRSS11D, 5X SYPRO orange dye, and 50 mM Tris pH 8.0 with 200 mM NaCl. Assays were performed in technical triplicate (*n=*3) and data are shown as mean +/− s.d. The ΔT_m_ data were curve-fitted for one-site EC_50_ in GraphPad Prism.

To probe differences between the TMPRSS2 and TMPRSS11D proteins that could impact nafamostat inhibition potency and inhibition *t*_1/2_, we used differential scanning fluorimetry (DSF) to measure ligand-induced shifts in protein T_m_s. We previously showed that 1 µM nafamostat induces a dasTMPRSS2 T_m_ shift (ΔT_m_) of 25.5±0.1°C^10^. Nafamostat induced dasTMPRSS11D ΔT_m_s from 0.10±0.05°C to 9.2 ±0.2°C for ligand concentrations spanning 0.1-100 µM (teal datapoints; Fig. 3d). When plotted on a semi-logarithmic scale, the ΔT_m_ values for dasTMPRSS11D induced by nafamostat reached saturation (teal datapoints; Fig. 3d). The data were curve-fitted to determine the half-maximal effective concentration (EC_50_) for thermal stabilization, which was 2.0±0.8 µM (teal trace; Fig. 3d). In contrast, 6-amidino-2-naphthol (dasTMPRSS11D K_i_ value of 71 µM; Extended Data Fig. 3a) induced a maximum dasTMPRSS11D ΔT_m_ of 4.67±0.06°C at a compound concentration of 10 mM, and ΔT_m_ values did not fully saturate (violet datapoints; Fig. 3d). When dasTMPRSS11D S368A was incubated with 0.01-100 µM nafamostat, no dasTMPRSS11D S368A ΔT_m_s were detected (violet; Extended Data Fig. 3b). In contrast, 6-amidino-2-naphthol induced a maximum eTMPRSS11D S368A ΔT_m_ of 1.7±0.2°C (teal; Extended Data Fig. 3b). These biophysical data confirm that nafamostat relies on the TMPRSS11D S368A residue to induce significant ΔT_m_ changes, likely through the formation of a 4-guanidino benzoate acyl-enzyme complex. However, nafamostat provided less thermal stabilization to TMPRSS11D compared to TMPRSS2. This difference may offer molecular insights into why the TMPRSS11D acyl-enzyme complex hydrolyzed more quickly than the TMPRSS2 complex. Overall, these kinetic and biophysical findings suggest that TMPRSS11D interacts with nafamostat like a high-affinity substrate (with a fast on-rate and slow off-rate) rather than as a potent, irreversible inhibitor of TMPRSS11D protease activity.

### The cleaved TMPRSS11D zymogen activation motif can bind to nearby molecules within a protein crystal lattice

To better understand TMPRSS11D’s esterase activity towards nafamostat, we set up cocrystallization experiments with dasTMPRSS11D preincubated with nafamostat to determine the structure of the acyl-enzyme complex. Protein crystals formed in two distinct precipitant conditions, but no electron density was observed attached to the S368 residue that was expected from the nafamostat co-structure determined for TMPRSS2^10^. Instead, the protease crystallized with the cleaved zymogen activation motif interacting with a neighboring protease molecule in the crystal lattice (magenta squiggle; Fig. 4a-b). The dasTMPRSS11D structure was solved at a resolution of 1.59 Å and the zymogen activation motif (DDDDK^186^–CO_2_^−^) was clearly resolved within the substrate binding cleft (Fig. 4c). We used this crystal form to next determine the structure of TMPRSS11D complexed with the more biologically relevant TMPRSS11D zymogen motif (LSEQR^186^–CO_2_^−^) by crystallizing eTMPRSS11D S368A and solving its structure at 1.90 Å resolution (Fig. 4d). Data collection and refinement statistics are summarized in Extended Data Table 2.

**Figure 4.**
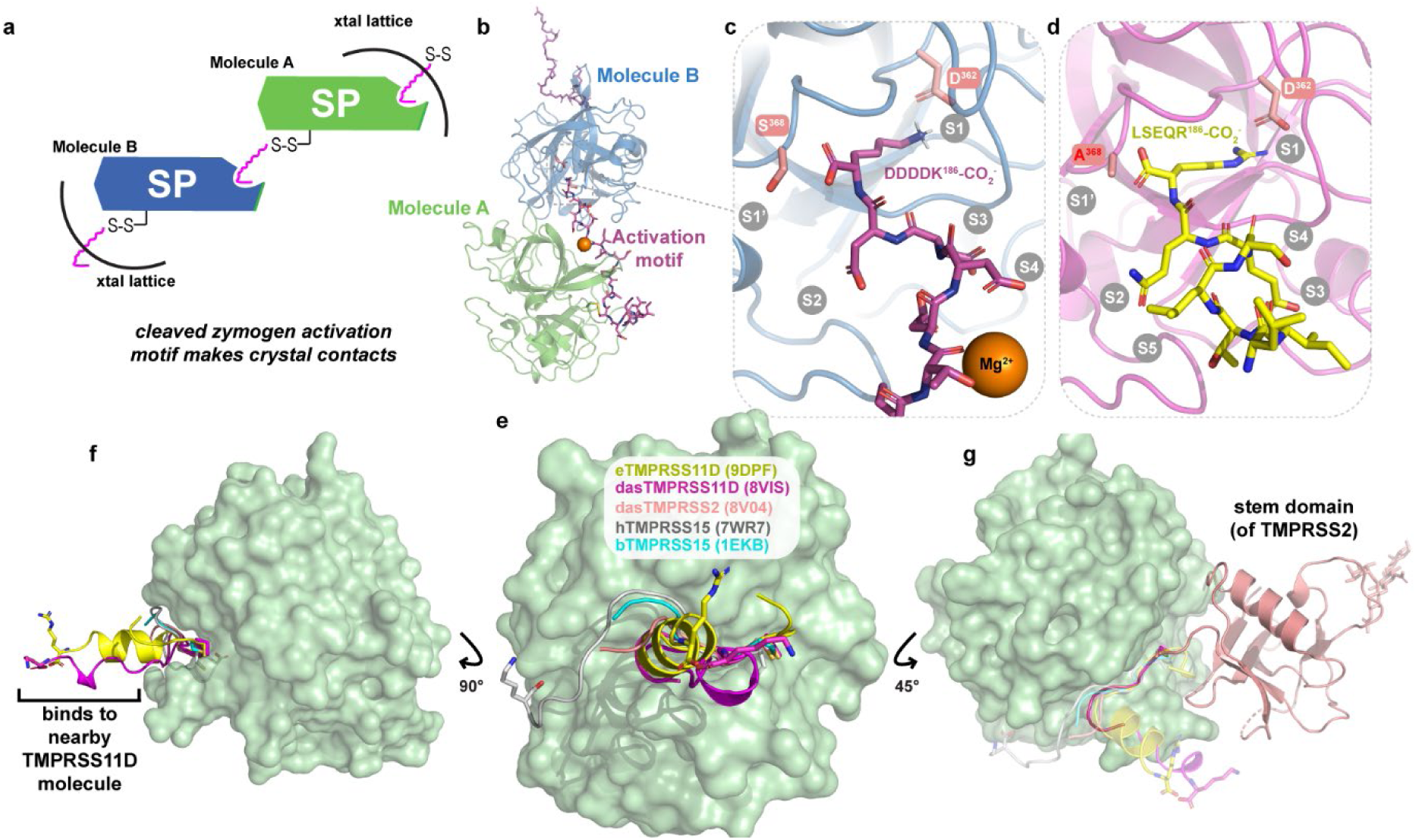
TMPRSS11D crystallizes by using intermolecular contacts with its own zymogen motif peptide occupying the substrate binding cleft. **a**, Schematic of the TMPRSS11D crystal lattice containing the TMPRSS11D serine protease (SP) domain and its cleaved zymogen activation motif (magenta squiggle). **b**, Cartoon representation of the dasTMPRSS11D crystal structure (PDB 8VIS). The cleaved zymogen activation motif (magenta sticks) of Molecule A interacts with the substrate binding cleft of Molecule B. **c**, Zoomed-in view of the dasTMPRSS11D active site occupied by the DDDDK^186^–CO_2_^−^ peptide. The TMPRSS11D catalytic Ser368 residue and the TMPRSS2 Subsite 1 (S1 residue) D362 are shown as salmon sticks. Additional TMPRSS11D subsites are denoted in gray text. **d**, Zoomed-in view of the eTMPRSS11D S368A active site occupied by the LSEQR^186^–CO_2_^−^ peptide (PDB 9DPF). **e**, Comparison of the cleaved zymogen activation motifs (attached through a disulfide bond) of the indicated TTSPs, with the SP domain of TMPRSS11D shown as a green surface. The terminal residues of the zymogen motifs of dasTMPRSS11D, eTMPRSS11D S368A, and human TMPRSS15 are shown as sticks. **f**, Side view of the zymogen activation motifs from (**e**). **g**, Side view of the back side of the TMPRSS11D SP domain. The stem domain of TMPRSS2 (salmon cartoon; PDB 7MEQ) is shown and is covalently attached to the SP domain of TMPRSS2 through an interdomain disulfide bond.

Only a few structures of active TTSPs contain any electron density for the disulfide-linked, cleaved zymogen activation motif. This is because TTSP proteins purified for crystallization are typically overexpressed as inclusion bodies in *Escherichia coli* and exclusively contain the TTSP SP domain^29–31^. Notable exceptions to this protein production and crystallization trend include TMPRSS1 (hepsin; PDB 1Z8G)^32^, TMPRSS2 (PDB 7MEQ)^10^, TMPRSS13 (PDB 6KD5)^33^, bovine enteropeptidase (PDB 1EKB)^34^, and human enteropeptidase (PDB 7WR7)^35^ which showed some electron density for their cleaved zymogen activation motifs. However, only a few zymogen motif amino acids were resolved for TMPRSS1, TMPRSS2, TMPRSS13, and bovine enteropeptidase at residues 5, 6, 6, and 7, respectively (Fig. 4e), while the cryo-EM structure of enteropeptidase (3.10 Å resolution) modelled the complete CGKKLAAQDITPK^186^–CO_2_^−^ zymogen activation motif. Interestingly, both the eTMPRSS11D S368A zymogen activation motif (yellow) and dasTMPRSS11D zymogen activation motif (magenta) adopted an α-helical structure oriented perpendicular to the face of the SP domain, whereas all other TTSP zymogen motifs were β-strand structures wrapped around the SP domain and/or the electronic density was poorly resolved around this region (Fig. 4e-f).

The TMPRSS11D SEA domain was not resolved in either TMPRSS11D crystal structure. To confirm that the domain was proteolytically cleaved, eTMPRSS11D S368A crystals were harvested, washed in buffer and subjected to SDS-PAGE separation and Coomassie blue staining (Extended Data Fig. 4a-c). Only a single protein band was detected that migrated at a molecular weight of ∼25 kDa which suggested that the TMPRSS11D SEA domain was not present in the protein crystal (Extended Data Fig. 4d). To predict where the SEA domain may interact with the TMPRSS11D SP domain, we superposed the TMPRSS11D crystal structure with the structure of TMPRSS2 which contained its active SP domain and SRCR domain linked by a disulfide bond (Fig. 4g). The SRCR domain of TMPRSS2, as well as other stem domains of TTSPs, supports the back face of the SP domain (salmon cartoon; Fig. 4g). This suggests that the SEA domain of TMPRSS11D may also be placed in that region prior to its proteolytic cleavage and shedding.

### The TMPRSS11D substrate binding cleft has unique and targetable features

TTSP crystal structures are valuable tools for atomic level understanding of the substrate binding cleft and the intermolecular interactions with both substrate or inhibitor ligands. In particular, peptides that contain Position 1 (P1) Arg/Lys residues^36^. Ligand residues C-terminal to P1 are termed P1’, P2’, etc., whereas residues N-terminal to P1 are termed P2, P3, etc. In general, the binding pockets responsible for recognizing each residue (S1, S1’, and S2, etc.) of a TTSP are well defined and predictably interact with ligand P1, P1’, and P2 residues (Fig. 5a). The P3 and P4 residues of a ligand are positioned farther from the catalytic triad and can adopt different conformations to interact with the S3 and S4 regions of the protease’s substrate-binding cleft. As a result, defining the S3 and S4 regions of a TTSP is somewhat ambiguous. This ambiguity became evident when the first structure of TMPRSS13 was solved, showing an inhibitor peptide making unexpected contacts between its P3 and P4 ligand residues with the canonical S3 and S4 subsites^33^. To explore targetable sites within the substrate binding cleft of TMPRSS11D, we compared all structurally validated subsites (with high resolution crystal structures) and predicted subsites (through a multiple sequence alignment) present in the TTSP family (Fig. 5b). The crystal structures of hepsin, TMPRSS13, enteropeptidase, and matriptase were determined in the presence of peptide inhibitors, allowing us to assign the protease amino acids comprising S1’, S1, S2, S3, and S4 and compare them on the surface of TMPRSS11D (Fig. 5c).

**Figure 5.**
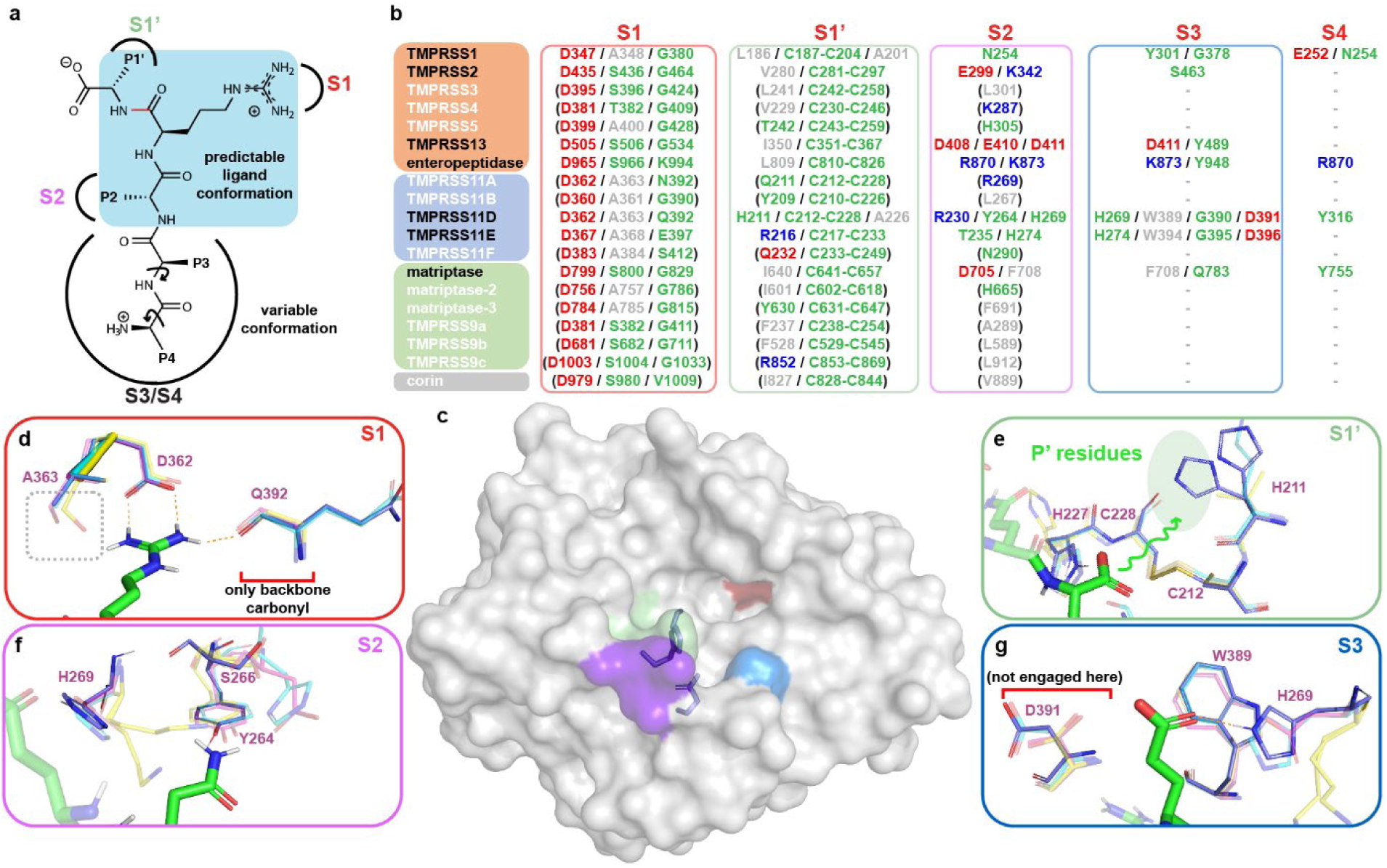
Structural comparison of the S1’, S1, S2 and S3 binding sites of TMPRSS11D, TMPRSS11E, TMPRSS2 and TMPRSS13. **a**, Schematic of a typical peptide ligand interacting with a TTSP, with the scissile peptide bond shown in red. The P1’, P1 and P2 ligand amino acids are highlighted in light blue and adopt predictable conformations to engage the S1’, S1, and S2 binding sites of the TTSP, respectively. **b**, Comparison of the amino acids at each protease subsite for every TTSP. TTSPs with experimental crystal structures are indicated in black text. Protease amino acids predicted to form salt bridges are indicated in red (electronegative) or blue (electropositive) whereas hydrophobic amino acids are indicated in gray text. Amino acids in green are predicted to participate in H-bonding with ligands. **c**, Surface representation of TMPRSS11D, with the S1’ (green), S1 (red), S2 (purple), and S3 (blue) subsites colored. The TMPRSS11D catalytic triad, S368-H227-D272, are shown as sticks. **d**, S1 of TMPRSS11D shown in stick representation. The R186 ligand residue (from PDB 9DPF) is shown as green sticks. H-bonds between the ligand and the TMPRSS11D S1 are shown as dashed orange lines. TMPRSS11D residues-purple TMPRSS11E (PDB 2OQ5)-teal, TMPRSS2 (PDB 8V04)-yellow, TMPRSS13 (PDB 6KD5)-pink. **e**, S1’ of TMPRSS11D. The carboxylate of R^186^–CO_2_^−^ is shown as green sticks and points towards S1’ (indicated by a green squiggle). **f-g**, S2 and S3 of TMPRSS11D.

The TMPRSS11D S1 is formed by D362 and Q392 that mediate salt bridges and H-bonds, respectively, with the P1 Arg (Fig. 5d). Unlike TMPRSS2 and TMPRSS13, the TMPRSS11D S1 subsite contains A363 instead of a Ser residue that typically H-bonds with the guanidine of P1 Arg residues. Furthermore, a Gly residue is typically present at the equivalent position of the TMPRSS11D Q392 residue for most other TTSPs, excepting enteropeptidase (Lys), TMPRSS11E (Glu), TMPRSS11A (Asn), and corin (Val; Fig. 5b). Thus, TMPRSS11D’s lack of the conserved Ser and Gly residues within S1 may explain why nafamostat and 6-amidino-2-naphthol did not induce as large of a ΔT_m_ for TMPRSS11D as for TMPRSS2. Furthermore, the fewer molecular contacts made by the nafamostat acyl-enzyme complex could lead to the rapid hydrolysis that is observed for TMPRSS11D but not TMPRSS2.

The S1’ of TMPRSS11D includes the conserved C212-C228 disulfide found in every TTSP (Fig. 5e). However, TMPRSS11D has a unique H211 residue whereas TMPRSS11E employed R216, TMPRSS13 used I350, and TMPRSS2 used V280, at their S1’. These subtle differences may confer some unique P1’ residue preference to TMPRSS11D compared to these other proteases. However, since the LSEQR^186^–CO_2_^−^ and DDDDK^186^–CO_2_^−^ peptides that were co-crystallized with TMPRSS11D did not contain P1’ residues, it is difficult to make conclusions about TMPRSS11D P1’ residue preferences.

The S2 of TMPRSS11D contains a unique combination of R230, Y264, and H269 which is distinct from every other TTSP (Fig. 5f). The H269 residue was also found as part of TMPRSS11D S3, which also contained W389, G390, and D391 (Fig. 5g). However, the TMPRSS11D S3 exactly matched the residues comprising the S3 of TMPRSS11E. In our TMPRSS11D crystal structures, no ligand residues contacted D391. The P4 ligand residue in the eTMPRSS11D S368A was mostly solvent exposed but made a single H-bond with the TMPRSS11D Y316 residue. Taken together, these structural data suggest that selective TMPRSS11D inhibitors may require specific contacts made with residues within TMPRSS11D S1’ and/or S2.

### A DYDDDDK-CO_2_^−^ peptide is a weak competitive inhibitor of dasTMPRSS11D

To confirm that the cleaved zymogen activation motif of dasTMPRSS11D, CGAGPDLITDDDDK^186^–CO_2_^−^, was not negatively impacting protease activity assays, we measured dasTMPRSS11D activity in the presence of high concentrations of a synthetic DYKDDDDK-CO_2_^−^ peptide (Extended Data Fig. 5a). Since a Mg^2+^ ion was observed interacting close to the substrate binding pocket in the dasTMRSS11D crystal structure, activity assays were performed in assay buffer containing 0-50 mM MgCl_2_ to evaluate if Mg^2+^ was facilitating TMPRSS11D autoinhibition with the DDDDK-CO2-zymogen activation motif. At a DYKDDDDK-CO_2_^−^ peptide concentration of 5 mM, dasTMPRSS11D peptidase activity was reduced by 75%. The MgCl_2_ concentration in the assay did not influence dasTMPRSS11D peptidase activity. Activity assays were also performed in various concentrations of NaCl to probe if protein-protein interactions between dasTMPRSS11D protease molecules were influencing peptidase activity (Extended Data Fig. 5b). TMPRSS11D peptidase activity was enhanced in assay buffer containing high salt concentrations (500 mM). These results suggest that while the DYKDDDDK-CO_2_^−^ peptide can inhibit dasTMPRSS11D activity, it is unlikely that the cleaved DDDDK^186^–CO_2_^−^, zymogen motif attached to dasTMPRS11D was substantially interfering with the catalytic activity of the enzyme, especially when assays were conducted with nanomolar concentrations of dasTMPRSS11D.

### Modelling the zymogen activation and shedding of the TMPRSS11D ectodomain

The TMPRSS11D SEA domain was not resolved in our TMPRSS11D crystal structure, motivating us to investigate modelled structures to develop a structure-informed understanding of full length TMPRSS11D zymogen activation and potential shedding from the cell surface. The AlphaFold2 TMPRSS11D structure (AF-O60235-F1) of the zymogen form of the protease modeled the zymogen activation motif in a similar orientation as that of the pro-matriptase structure (Fig. 6a; Extended Data Fig. 6a). As expected, the SEA domain of the Alphafold2 TMPRSS11D structure superposed closely with the 1.92 Å resolution crystal structure of the SEA domain of mouse TMPRSS11D (PDB 2E7V; Extended Data Fig. 6b). Furthermore, the placement of the SEA domain relative to the TMPRSS11D SP domain was similar to the TMPRSS2 crystal structure containing its SP and SRCR domains, but the SEA domain was larger and extended in the opposite direction of the SRCR domain (Extended Data Fig. 6c). Thus, the AlphaFold2 TMPRSS11D structure serves as a plausible prediction of the zymogen form of the full-length protease.

**Figure 6.**
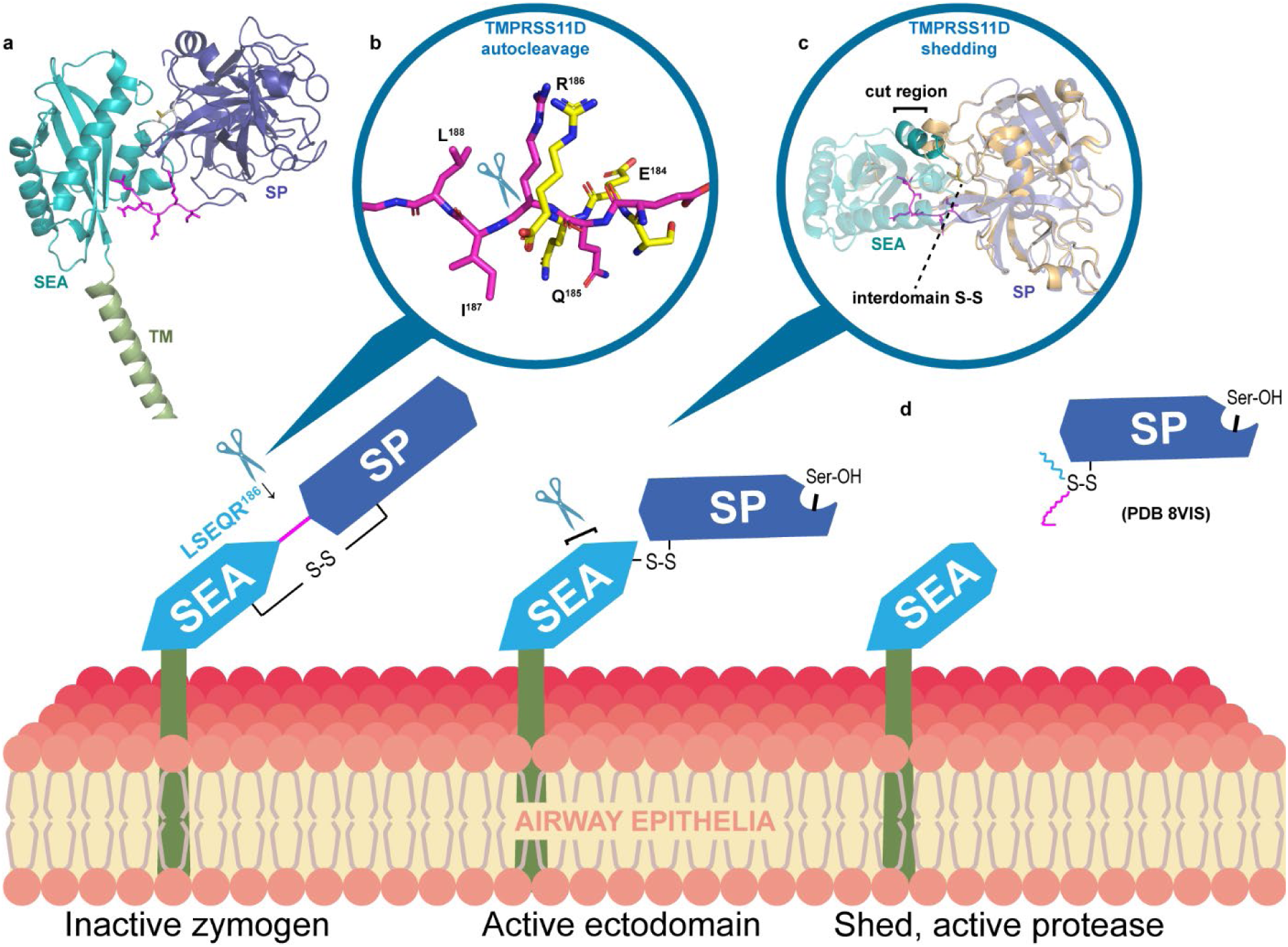
A proposed model of TMPRSS11D autocleavage activation and shedding from the cell surface. **a**, AlphaFold 2.0 model of full-length human TMPRSS11D (AF-O60235-F1). The transmembrane (TM) domain (green), the Sea urchin, Enteropeptidase and Agrin (SEA) domain (teal), and the serine protease (SP) domain (purple) are shown in cartoon representation. The TMPRSS11D zymogen activation motif is shown in magenta sticks. **b**, Superposed model of the AlphaFold 2.0 TMPRSS11D protein structure (magenta sticks) and the cleaved TMPRSS11D zymogen activation motif (PDB 9DPF; yellow sticks). The TMPRSS11D cleavage site spanning the R186-I187 peptide bond is denoted with a scissor graphic. **c**, Cartoon representation of the TMPRSS11D cleavage event in the SEA domain leading to protease shedding from the cell surface. The SP domains of the AlphaFold 2.0 TMPRSS11D structure (purple) and the dasTMPRSS11D crystal structure (cartoon) were superposed. The suspected SEA domain cleavage site is indicated with a black bar. **d**, Graphical representation of the shed TMPRSS11D SP domain.

After the full-length TMPRSS11D zymogen reaches the cell surface, it is possible that an active TMPRSS11D protease molecule (or another trypsin-like serine protease) recognizes the zymogen motif as a substrate and cleaves the R186-I187 peptide bond. Accordingly, we investigated if the conformation of the zymogen motif was appropriate for cleavage by TMPRSS11D (Fig. 6b). We superposed the cleaved zymogen motif of the eTMPRSS11D S368A structure (LSEQR^186^–CO_2_^−^; yellow sticks; Fig. 6b) upon the intact, exposed zymogen motif of the AlphaFold2 TMPRSS11D structure (magenta sticks; Fig. 6b). The conformations of the ligands were similar and indicated that an active TMPRSS11D molecule could potentially bind and cleave this exposed zymogen motif depicted in the AlphaFold2 structure. After the TMPRSS11D molecule has undergone zymogen activation, it can exert its proteolytic function at the cell surface. Our model also is consistent with subsequent cleavage by TMPRSS11D of a residue near the C-terminus of the SEA domain (bolded teal cartoon; Fig. 6c) thereby enabling shedding of the active TMPRSS11D SP domain from the cell surface (Fig. 6d).

## DISCUSSION

The zymogen activation step of TTSPs is a critical aspect of their biology and determines when the TTSP harbors its matured proteolytic activity. Each TTSP family member has a unique sequence of amino acids within their zymogen activation motifs. The biochemical and structural data presented here showed that TMPRSS11D and TMPRSS2 recognized their own zymogen activation motifs, allowing them to turn on their own proteolytic activity. To facilitate their analysis, we replaced their zymogen activation motifs with DDDDK; this delayed their zymogen activation and enabled protein overexpression, biochemical assays, and protein crystallization. We confirmed that the DDDDK sequence does not interfere with protease activity assays for high-throughput drug screening applications. Furthermore, these proteases efficiently cleave the SARS-CoV-2 S protein in functional protease activity assays.

Nafamostat mesylate has been proposed as an antiviral agent disabling TTSPs and has repeatedly been used in cell studies to block TTSP-mediated viral entry at the cell surface^3,5,37^. Our kinetic data indicated that TMPRSS11D interacted with nafamostat as a substrate, rapidly forming an acyl-enzyme complex and then hydrolyzing the ester to convert it to the product molecules 4-guanidino benzoic acid and 6-amidino-2-naphthol (Fig. 3). 6-amidino-2-naphthol inhibited TMPRSS11D, but with much weaker potency than its parent molecule nafamostat. Thus, TMPRSS11D may not be disabled by nafamostat (and other related esters such as camostat) over a prolonged period. TMPRSS11D’s esterase activity suggested that ester-based compounds may not be effective as antivirals for cells expressing this enzyme. When developing ester drugs targeting TTSPs, the rate at which ester compounds are broken down should be kinetically characterized^10,38^. Additionally, an ester that is a potent inhibitor of a particular TTSP target could be rapidly turned over by another co-expressed TTSP and rapidly eliminate the compound.

The TMPRSS11D crystal structure provided some molecular insights into the reversibility of nafamostat inhibition. The TMPRSS11D S1 lacks an H-bond donor relative to TMPRSS2’s S1, and nafamostat cannot thermally stabilize (measured through nafamostat-induced ΔT_m_s) TMPRSS11D at the same magnitude as TMPRSS2^10^. All other HAT/DESC TTSP subfamily members lack this H-bond donor residue, as they all contain Ala residues at this position in their S1 (Fig. 5b). Other electrostatic and steric features of the substrate binding cleft may promote hydrolysis of the acyl-enzyme complex. Overall, the protease activity of the HAT/DESC subfamily of TTSPs may not be as effectively disabled by nafamostat as other TTSP subfamilies.

The dasTMPRSS11D crystallization trials were initially performed in the presence of high concentrations of nafamostat or 6-amidino-2-naphthol. Protein crystals formed in the presence of each ligand, but the final crystal structures contained no electronic density matching the ligands. Instead, the zymogen activation motif of dasTMPRSS11D, DDDDK^186^–CO_2_^−^, was found occupying the substrate binding cleft of a nearby TMPRSS11D molecule within the crystal lattice. We repeated this crystallization strategy for TMPRSS11D with its native zymogen activation motif, LSEQR^186^–CO_2_^−^, and determined the first crystal structure of a TTSP interacting with its own (natural) zymogen activation motif. This structure explained why dasTMPRSS11D was capable of cleaving and activating eTMPRSS11D S368A and explained why TMPRSS11D can autoactivate in cells^39^. Using the dasTMPRSS11D and eTMPRSS11D S368A crystal structures, we were able to map the S1’-S4 subsites of TMPRSS11D which could serve as targetable regions of the TMPRSS11D substrate binding pocket in drug development campaigns using peptidomimetic or small molecule inhibitors. The S1’ of TMPRSS11D is distinct with its H211 residue and could enable unique stacking interactions with an aromatic or hydrophobic P1’ residue. The S2 of TMPRSS11D contains a unique combination of R230 and H269 which is distinct from every other TTSP. The S3 and S4 sites of TMPRSS11D are distinct from every other TTSP (except TMPRSS11E) and feature a combined H269 and D391 cluster. As described earlier, the conformation of the peptide ligand at the P3 and P4 positions determines what protease subsites they will interact with. Positional scanning libraries may provide general insights into protease preferences, but the crystal structures of the protease bound to preferred peptides may better inform what protease amino acid subsites the ligand is interacting with.

The protease crystallization strategy identified here may be generally applicable to TTSPs. Notably, other TTSPs have been structurally characterized with their disulfide-linked zymogen activation motifs placed in the same region as our crystal structures at the back of the TTSP SP domain (Fig. 4). However, the TTSP stem domains may sterically block the crystallization tag from interacting with a neighboring protease molecule in the crystal lattice. To test this, we overexpressed and purified dasTMPRSS11D with a S368A mutation, and carefully activated the protease through treatment with enteropeptidase (Extended Data Fig. 7). The protease was fully activated by enteropeptidase treatment and had the same DDDDK-CO_2_^−^ peptide available to act as a crystallization tag as the original dasTMPRSS11D crystal structure. However, the SDS-PAGE gel migration patterns of this sample suggested that the protein contained the SEA domain (Extended Data Fig. 7b) so the crystallization experiment contained the complete TMPRSS11D ectodomain. No protein crystals formed under the same precipitant screening conditions as dasTMPRSS11D and eTMPRSS11D S368A. Interestingly, the SEA domain has previously been reported to impede both protein crystallization and structure determination by cryo-EM^35^. Thus, to reapply this crystallization strategy to other TTSPs, the stem domains of the protease may need to be removed to enable access to the crystallization tag by neighboring protease molecules, with recombinant protein construct designs informed by the TMPRSS11D crystal structures here.

The biochemical and structural data outlined here have provided an improved understanding of TMPRSS11D and TMPRSS2 zymogen cleavage activation and explained how these proteases may be capable of autocleavage activation in human cells. Thus, TMPRSS2 and TMPRSS11D are two TTSPs that rapidly autoactivate *in vitro* and are efficient drivers of SARS-CoV-2 infection *in vitro* and *in vivo*^7^. TTSP autocleavage activation may therefore be an indicator of a protease’s ability to drive respiratory virus infections, and future work may help define how TTSP zymogen activation influences viral pathobiology and help prioritize human protease targets for antiviral development.

## CONCLUSION

TTSPs are exceptionally valuable but difficult drug targets owing to their high structural conservation with other trypsin-like serine proteases important for human biology. Selectively targeting them requires key biochemical and structural insights, and our recombinant protein production and crystallization tag strategy developed here may enable the TTSP family to be rapidly interrogated with peptide-bound ligands that uncover regions for selective pharmacological targeting. We note that the ester compound nafamostat is rapidly cleaved by TMPRSS11D and potentially other TTSPs, potentially explaining its lack of efficacy as an antiviral agent. These data also provide key structural understandings of protease zymogen activation.

## METHODS

### Construct design and cloning

TMPRSS11D cDNA (nucleotide accession # BC125196) was purchased from Transomic and constructs encoding the soluble TMPRSS11D ectodomain spanning residues 44-418 were subcloned into the pFHMSP-LIC C or pFHMSPN-avi-TEV-LIC baculovirus donor vectors. All constructs contained a N-terminal honeybee melittin signal sequence peptide. For pFHMSP-LIC C, proteins had a C-terminal His8 tag. For pFHMSPN-avi-TEV-LIC, proteins had a N-terminal His6 tag followed by an Avi tag for biotinylation, then a TEV cleavage site. All mutagenesis primers are listed in Supplementary Table 1 and mutagenesis was achieved using the LIC method. For the engineered active TMPRSS11D construct (dasTMPRSS11D), L182D/S183D/E184D/Q185D/R186K mutations were implemented. The donor vectors containing the engineered TMPRSS11D gene were transformed into *Escherichia coli* DH10Bac cells (Thermo Fisher; Cat# 10361012) to generate recombinant viral bacmid DNA. Sf9 cells were transfected with Bacmid DNA using JetPrime transfection reagents (PolyPlus Transfection Inc.; Cat# 114-01) according to the manufacturer’s instructions, and recombinant baculovirus particles were obtained and amplified from P1 to P2 viral stocks. P1 viral stocks were used for protein test expression studies with suspension culture of baculovirus infected insect cells. For scaled-up productions of TMPRSS11D protein, recombinant P2 viruses were used.

The SARS-CoV-2 Spike ectodomain HexaPro construct was a gift from J. McLellan, and the S1/S2 site was restored (GSAS685->RRAR; HexaFurin construct) through site-directed mutagenesis as previously described^10^.

### Baculovirus mediated TMPRSS11D protein production in Sf9 insect cells

Sf9 cells were grown in I-Max Insect Medium (Wisent Biocenter; Cat# 301-045-LL) to a density of 4×10^6^ cells/mL and infected with 20 mL/L of suspension culture of baculovirus infected insect cells prior to incubation on an orbital shaker (145 rpm, 26 °C).

### Recombinant TMPRSS11D protein purifications

TMPRSS11D ectodomain proteins were produced through secreted expression and purified using a similar protocol to TMPRSS2^10^. Cell culture medium containing the final secreted protein products AA-[TMPRSS11D (44-418)]-EFVEHHHHHHH (for dasTMPRSS11D) or AAPEMHHHHHHEFMSGLNDIFEAQKIEWHEGSAGGSGENLYFQG-[TMPRSS11D (44-418) (for eTMPRSS11D S368A and dasTMPRSS11D S368A) were collected by centrifugation (20 min, 10 °C, 6,000 x g) 4-5 days post-infection when cell viability dropped to 55 - 60%. Media was adjusted to pH 7.4 by addition of concentrated PBS stock, then supplemented with 15 mL/L settled Ni^2+^-NTA resin (Qiagen) at a scale of 3 L and distributed at a scale of 1.5 L to 2.8L glass flasks. Flasks were shaken for 1 hour at 16 °C (110 rpm), then bead-media mixtures were transferred to 0.5 L gravity flow columns (Bio-Rad). Beads were washed with 3 column volumes (CVs) PBS prior to elution with 1.5 resin bed volumes of Elution Buffer (PBS supplemented with 250 mM imidazole). Crude protein was concentrated using 10 kDa MWCO Amicon filters and washed into PBS to remove excess imidazole. Protein samples were prepared for SDS-PAGE with either reducing (5 mM 2-mercaptoethanol) Laemelli dye and thermally denatured for 5 minutes, or nonreducing Laemelli dye and were not thermally denatured. Concentrated Ni^2+^-NTA IMAC elution samples were passed through a 0.22 µm syringe filter and injected to a Superdex75 gel filtration column pre-equilibrated with Size-Exclusion Chromatography (SEC) Buffer (50 mM Tris pH 8.0, 200 mM NaCl). SEC fractions containing TMPRSS11D proteins were pooled, concentrated to 5 mg/mL, then flash frozen in liquid nitrogen and stored as aliquots at −80 °C in advance of assays.

### Crystallization and data collection for dasTMPRSS11D and eTMPRSS11D S368A

SEC-purified dasTMPRSS11D protein (3 mg/mL) was incubated with excess nafamostat (5:1 compound:protease) or excess 6-amidino-2-naphthol (100:1 compound:protease) at 4 °C for 15 min, then concentrated to 30 mg/mL using a 10 kDa MWCO Amicon filter and centrifuged (14,000 rpm, 10 min, 4 °C). SEC-purified eTMPRSS11D S368A protein (at 20 mg/mL) was similarly prepared for crystallization trials but was not incubated with nafamostat or 6-amidino-2-naphthol. Protein samples were subjected to automated screening at 18 °C in 96-well Intelliplates (Art Robin) using the Phoenix protein crystallization dispenser (Art Robbins). Protein was dispensed as 0.5 µL sitting drops and mixed 1:1 with precipitant. The RedWing and SGC precipitant screens were tested with a large crystal obtained for the nafamostat:dasTMPRSS11D sample with precipitant solution containing 20% PEG1500, 0.2 M MgCl_2_, and 0.1 M HEPES pH 7.5. The 6-amidino-2-naphthol:dasTMPRSS11D sample produced crystals with precipitant condition containing 20% PEG3350 and 0.2 M Mg(NO_3_)_2_. For eTMPRSS11D S368A, protein crystals were obtained as sitting drops grown over precipitant solution containing 0.5 M Mg formate and 0.1 M Bis-Tris pH 6.5. To acquire diffraction-quality crystals, drops were reset as 2 µL hanging drops on glass slides. All diffraction-quality protein crystals were cryo-protected with reservoir solution containing 10% (*v*/*v*) ethylene glycol and cryo-cooled in liquid nitrogen. X-ray diffraction data were collected on the CMCF-ID beamline at the Canadian Light Source with a Dectris Eiger X 16M detector.

### Solving the TMPRSS11D crystal structures

X-ray diffraction data were processed with HKL-3000^40^. Initial phases for the dasTMPRSS11D crystal structure were obtained by molecular replacement using Phaser MR^41^ with the catalytic chain of TMPRSS11E (PDB 2OQ5) as a search model. The dasTMPRSS11D crystal space group was P4_3_2_1_2 and 4 TMPRSS11D SP domain protein molecules related by translational non-crystallographic symmetry were observed in the asymmetric unit. Model building was performed in COOT and refined with REFMAC5.8.0352^42^. The dasTMPRSS11D stem chain residues 166-186 were manually built into electron density within the TMPRSS11D substrate binding cleft. For eTMPRSS11D S368A, the crystal space group was P2_1_2_1_2_1_ and the structure was solved using molecular replacement with the dasTMPRSS11D crystal structure. Two TMPRSS11D SP domain protein molecules were found in the asymmetric unit. Both structures were validated by Molprobity^43^. The dasTMPRSS11D crystal structure (1.59 Å) and eTMPRSS11D S368A crystal structure (1.90 Å) were uploaded to the PDB under accession codes 8VIS and 9DPF, respectively.

### Protein sequence alignments and structure superpositions

The FASTA sequences of human TTSP family members were accessed through Uniprot (isoform 1) and aligned using Clustal Omega^44^ and annotated with ESPript v.3.0^45^. Protein structures were accessed from the PDB and superposed using PyMOL v2.5.7. Protein structure figures were prepared in PyMOL v2.5.7.

### TMPRSS11D peptidase assays and IC_50_ determination

dasTMPRSS11D peptidase assays with fluorogenic Boc-Gln-Ala-Arg-AMC substrate (Bachem Cat # 4017019.0025) were carried out using a similar protocol to dasTMPRSS2^10^. Assays were carried out at a scale of 50 µL in Greiner Black 384-well microplates, with AMC fluorescence monitored on a Molecular Devices FlexSation 3 plate reader (SoftMax Pro v5.4.6 software package) at excitation:emission of 360nm:480nm or on a BioTek Synergy H1 plate reader (Gen5 v3.03 software package) at 341nm:441nm where indicated. TMPRSS11D half-maximal inhibitory concentration (IC_50_) assays contained a final concentration of 0.2 nM dasTMPRSS11D with buffer containing 50 mM Tris pH 8.0, 250 mM NaCl, and 6mM CaCl_2_.

Nafamostat mesylate (MedChemExpress Cat# HY-B0190A) and 6-amidino-2-naphthol methanesulfonate (TCI Cat# A1193) were prepared as fresh DMSO stocks immediately prior to inhibition assays. To determine dasTMPRSS11D IC_50_s, inhibitor titrations prepared in DMSO were transferred to wells containing dasTMPRSS11D enzyme and incubated for 10 min. Boc-QAR-AMC substrate was transferred to wells (100 µM final concentration) and plates immediately read for AMC fluorescence. Initial reaction velocity slopes were tabulated across the first 0-120 seconds of the assay and normalized (as a percentage) relative to uninhibited enzyme. Inhibition data was curve-fitted for Absolute IC_50_ in Graphpad Prism and IC_50_ values were reported as mean values +/− standard deviation.

### TMPRSS11D inhibitor kinetic parameter determination

The dasTMPRSS11D covalent inactivation parameter *k*_inact_/K_I_ and noncovalent, competitive inhibition constant (K_i_) were determined using inhibitor coaddition assays described previously^10^ and reaction progress curves were curve-fitted using DynaFit 4.0. Stocks of Boc-QAR-AMC substrate with various concentrations of nafamostat (final concentrations of 1600, 800, 400, 200, 100, and 0.8 nM) or 6-amidino-2-naphthol (final concentrations of 500, 250, 125, 62.5, 31.3, 16, and 1 µM) were transferred to wells containing dasTMPRSS11D enzyme (3 nM final) using an Agilent Bravo liquid-liquid transfer device and AMC fluorescence was immediately read. Reaction progress was monitored for 30 min. The simplified Boc-QAR-AMC substrate conversion rate, *k*_sub_, was determined by curve fitting reaction progress curves in the presence of varying concentrations of dasTMPRSS11D enzyme using **Model 1** (Supplementary Fig. 1a). To determine the dasTMPRSS11D K_i_ for 6-amidino-2-naphthol, reaction progress curves were fit to **Model 2** and the dasTMPRSS11D *k*_inact_/K_I_ for nafamostat was determined using **Model 3** (Supplementary Fig. 1b-c).

The nafamostat inhibition *t*_1/2_ value was determined by pre-incubating 8 nM dasTMPRSS11D with various concentrations of nafamostat (1000, 500, and 250 nM) for 3 minutes prior to transfer to wells containing Boc-QAR-AMC substrate. Reaction progress was monitored for 3 hours and the data curve-fitted to determine the nafamostat hydrolysis rate (*k*_hydrolysis_) and nafamostat inhibition *t*_1/2_ value using **Model 4** (Supplementary Fig. 1d).

### SARS-CoV-2 S protein cleavage inhibition assays

The SARS-CoV-2 S protein construct HexaFurin was prepared as a substrate for dasTMPRSS11D as previously demonstrated for dasTMPRSS2^10^. dasTMPRSS11D was pre-incubated with 100-0.01 µM nafamostat (log_10_ dilution series) or 1000-62.5 µM 6-amidino-2-naphthol (log_2_ dilution series) prior to transferring the enzyme-inhibitor mixtures to 5 µg S protein substrate. S protein cleavage assays were carried out over 20 min at room temperature before assays were quenched through addition of 4X Laemmeli buffer. SDS-PAGE samples were thermally denatured (5 min at 95 °C) and approximately 4 µg S protein were loaded per well. After gel separation, protein bands were visualized by Coomassie blue staining.

### DSF for inhibitor-induced TMPRSS11D ΔT_m_s

TMPRSS11D protein T_m_s and ligand-induced ΔT_m_s were measured using SYPRO Orange dye (Life Technologies, catalog no. S-6650) and SYPRO fluorescence at 470 and 510 nm excitation and emission, respectively, using the Light Cycler 480 II (Roche Applied Science). Samples were prepared in technical triplicate in 384-well plates (Axygen; catalog nos. PCR-384-C; UC500) at a final volume of 20 µL. Wells contained 2 µg TMPRSS11D protein, 10% (*v*/*v*) ligand (or DMSO control) and 5× SYPRO Orange. Thermal melt curves were generated across a 25 to 95 °C gradient at a heating rate of 2 °C/min. TMPRSS11D T_m_ values +/− s.d. were calculated using the dRFU method with the DSFworld application^46^. Ligand-induced dasTMPRSS11D or dasTMPRSS11D S368A ΔT_m_s relative to DMSO control were calculated for each concentration of nafamostat or 6-amidino-2-naphthol and plotted using Graphpad Prism.

### Data exclusion and statistics

The two-sided Grubbs’ test was used to determine and exclude single outliers present in sample data performed in triplicate or quadruplet. Single datapoint outliers were identified in Fig. 2e (quadruplet samples, 1000, 10, and 0.001 µM 6-amidino-2-naphthol), Fig. 3d (triplicate samples, 0 µM nafamostat), Extended Data Fig. 2a (*n=*6 replicates, 1000 µM Boc-QAR-AMC initial reaction velocity), and Extended Data Fig. 5b (n=2 replicates, 0 µM DYKDDDDK and 0 mM MgCl_2_ sample). Within the supplied source data documents, these exclusions are denoted with blue text and asterisks.

## DATA AVAILABILITY

The coordinates and structures of dasTMPRSS11D complexed with a DDDDK peptide and eTMPRSS11D S368A complexed with a LSEQR peptide have been deposited to the PDB with accession numbers 8VIS (released January 31, 2024) and 9DPF (released October 16, 2024), respectively. The engineered dasTMPRSS11D and eTMPRSS11D S368A protein expression constructs are available on Addgene (Plasmid nos. 220881 and 220882, respectively). Zymogen matriptase, active matriptase, TMPRSS2, bovine enteropeptidase, human enteropeptidase, TMPRSS11E, and TMPRSS13 structures were accessed using PDB IDs 5LY0, 4JYT, 8V04, 1EKB, 7WR7, 2OQ5, and 6KD5, respectively.

## AUTHOR CONTRIBUTIONS

C.H.A., B.J.F., G.B.M. and F.B. provided project supervision; B.J.F conceived the project; B.J.F. and R.P.W. designed the experiments; Y.L. cloned TMPRSS2 and TMPRSS11D protein constructs for expression; A.S., R.P.W., O.I., Y-Y.L., and Z.H. produced TMPRSS2 and TMPRSS11D proteins in insect cells; B.J.F., O.I., and J.L. purified recombinant proteins; B.J.F. crystallized TMPRSS11D and B.J.F. and A.D. collected diffraction data; B.J.F, A.D., and T.MG.K. solved the TMPRSS11D crystal structures; B.J.F. performed bioinformatic and structural analyses; B.J.F., R.P.W. and J.L. performed fluorogenic peptidase activity and inhibitor potency assays and analyzed kinetics; B.J.F. and O.I. performed gel-based S protein digestion assays; B.J.F., O.I., and J.L. performed DSF assays; B.J.F. prepared figures; B.J.F., R.P.W., C.H.A., G.B.M., and A.M.E. wrote the manuscript.

## ACKNOWLEDGEMENTS

This work was supported by BC Leadership Chair in Functional Cancer Imaging to FB, and a Killam Doctoral Fellowship and a Mitacs Elevate Postdoctoral Fellowship to BF. This work is based upon research conducted using beamline CMCF-ID at the Canadian Light Source, a national research facility of the University of Saskatchewan, which is supported by the Canada Foundation for Innovation (CFI), the Natural Sciences and Engineering Research Council (NSERC), the National Research Council (NRC), the Canadian Institutes of Health Research (CIHR), the Government of Saskatchewan, and the University of Saskatchewan.. The Structural Genomics Consortium is a registered charity (no: 1097737) that receives funds from AbbVie, Bayer AG, Boehringer Ingelheim, Genentech, Genome Canada through Ontario Genomics Institute [OGI-196], the EU and EFPIA through the Innovative Medicines Initiative 2 Joint Undertaking [EUbOPEN grant 875510], Janssen, Merck KGaA (aka EMD in Canada and US), Pfizer, Takeda and the Wellcome Trust [106169/ZZ14/Z].

## EXTENDED DATA

**Extended Data Table 1.**
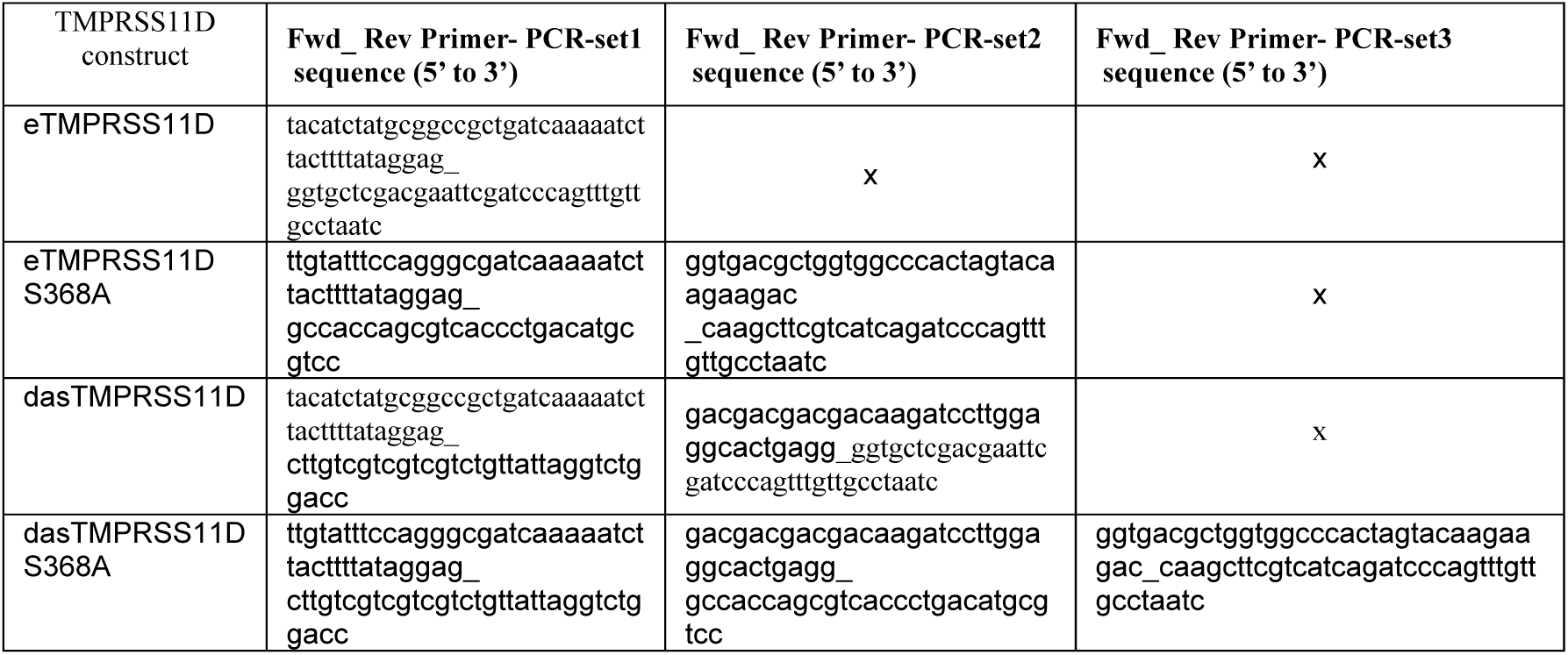

**Extended Data Table 2.** Data collection and refinement statistics for TMPRSS11D protein crystal structures.

|  | dasTMPRSS11D<br>(PDB 8VIS) | eTMPRSS11D S368A<br>(PDB 9DPF) |
| --- | --- | --- |
| **Data collection** |  |  |
| Space group | P4<sub>3</sub>2<sub>1</sub>2 | P2<sub>1</sub>2<sub>1</sub>2<sub>1</sub> |
| Cell dimensions |  |  |
| *a*, *b*, *c* (Å) | 119.68, 119.68, 134.29 | 73.03, 76.64, 88.37 |
| $\alpha$, $\beta$, $\gamma$ (°) | 90.00, 90.00, 90.00 | 90.00, 90.00, 90.00 |
| Resolution (Å) | 1.59(49.77-1.59) * | 1.90(40.0-1.90)* |
| *R*<sub>merge</sub> | 0.23 | 0.22 |
| *I* / $\sigma$ *I* | 1.57(1.59) | 1.60 (1.91) |
| Completeness (%) | 100.0 (49.77-1.59) | 99.2 (38.31-1.90) |
| Redundancy | 4.8 (4.3) | 11.4 (9.4) |
| **Refinement** |  |  |
| Resolution (Å) | 1.59 (49.77-1.59) | 1.90 (38.31-1.90) |
| No. reflections | 129049 | 37404 |
| *R*<sub>work</sub> / *R*<sub>free</sub> | 0.155 / 0.187 | 0.202 / 0.257 |
| No. atoms |  |  |
| Protein | 7732 | 3780 |
| Ligand/ion | 28 | 8 |
| Water | 1149 | 265 |
| *B*-factors | 17.0 | 34.0 |
| Protein | 14 | 32 |
| Ligand/ion | 20 | 59 |
| Water | 46.3 | 42.4 |
| R.m.s. deviations |  |  |
| Bond lengths (Å) | 0.55 | 0.47 |
| Bond angles (°) | 0.89 | 0.77 |

\*Each structure was solved using diffraction data from a single crystal. \*\*Values in parentheses are for highest-resolution shell.

**Extended Data Figure 1.**
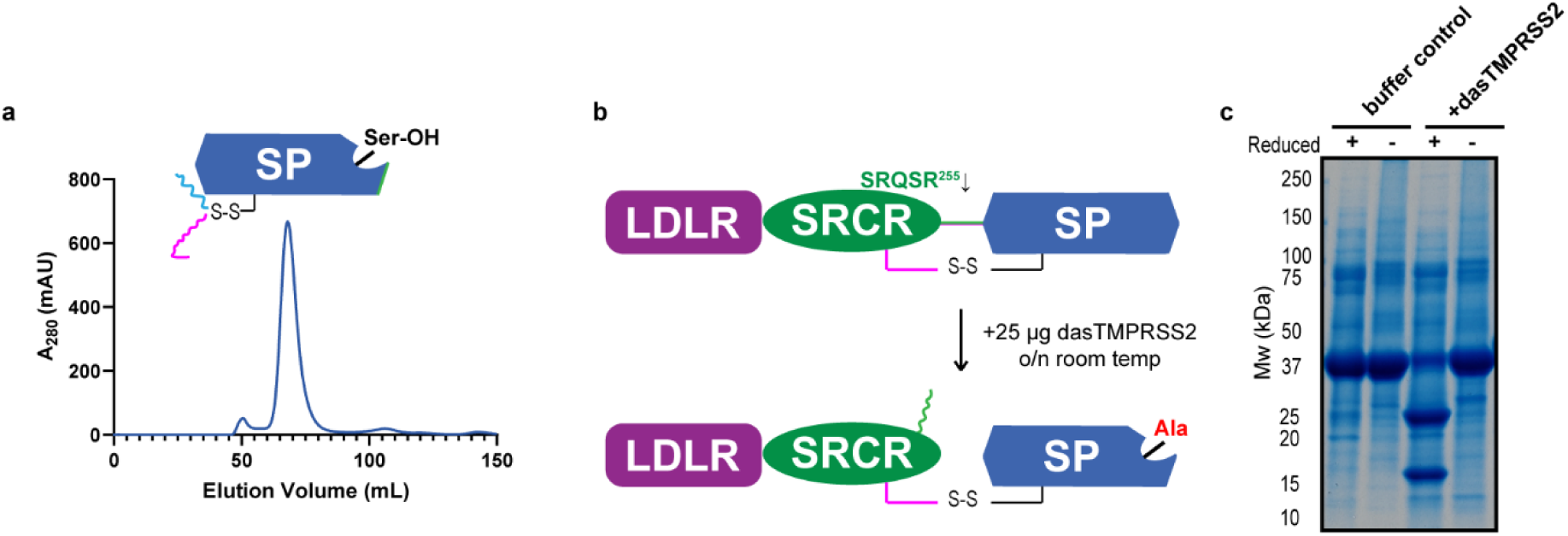
Purification of dasTMPRSS11D and demonstration that TMPRSS2 cleaves its own SRQSR255 zymogen activation motif. **a,** Size-exclusion chromatography purification of active dasTMPRSS11D. Approximately 18 milligrams of crude, active dasTMPRSS11D protein was loaded to a HiLoad 16/60 SuperDex 75 gel filtration column (4 °C, 1 mL/min, 50 mM Tris pH 8.0, 200 mM NaCl). Protein content was monitored by A_280_. **b,** Schematic of a soluble TMPRSS2 Ser441Ala protein construct before and after treatment with a trace amount of active dasTMPRSS2 protease. **c**, SDS-PAGE analysis of (**b**) for TMPRSS2 Ser441Ala protein incubated overnight with buffer control or 25 µg active dasTMPRSS2 protease.

**Extended Data Figure 2.**
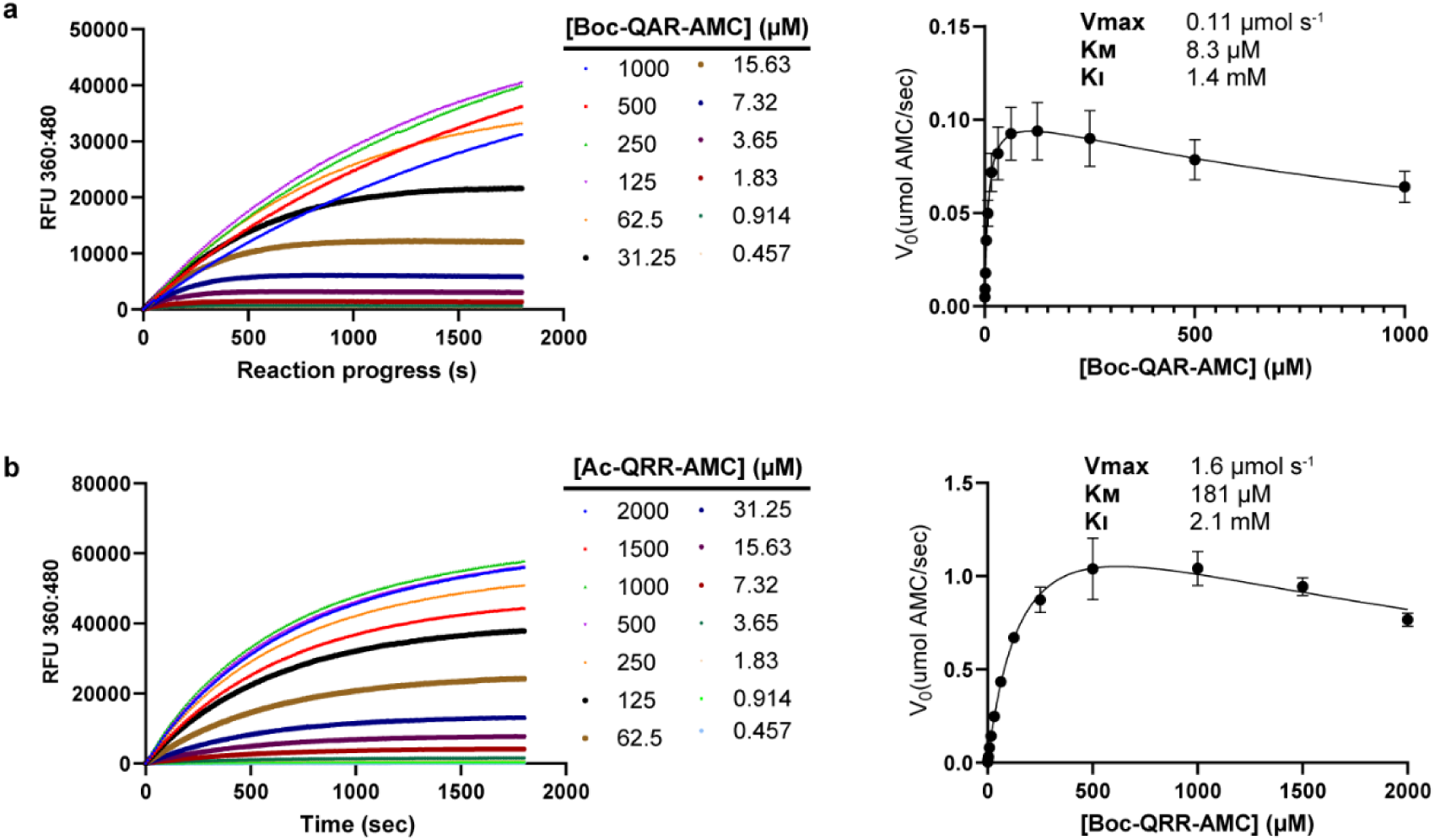
Kinetic characterization of fluorogenic peptide substrates with dasTMPRSS11D. **a**, (Left) Reaction progress curves of 3 nM dasTMPRSS11D protease with the indicated concentration of Boc-QAR-AMC peptide substrate, monitored by fluorescence (360:480). (Right) Michaelis-Menten plot of dasTMPRSS11D with Boc-QAR-AMC substrate. Initial reaction velocity plots were determined by taking the slope across the first 60 seconds of the reaction. Michaelis-Menten curves were fit in Graphpad Prism with a substrate inhibition parameter, producing the indicated values for maximum reaction velocity (V_max_), Michaelis constant (K_m_) and substrate apparent inhibition constant (K_i_). **b**, Reaction progress curves and Michaelis-Menten plot of dasTMPRSS11D with Ac-QRR-AMC substrate.

**Extended Data Figure 3.**
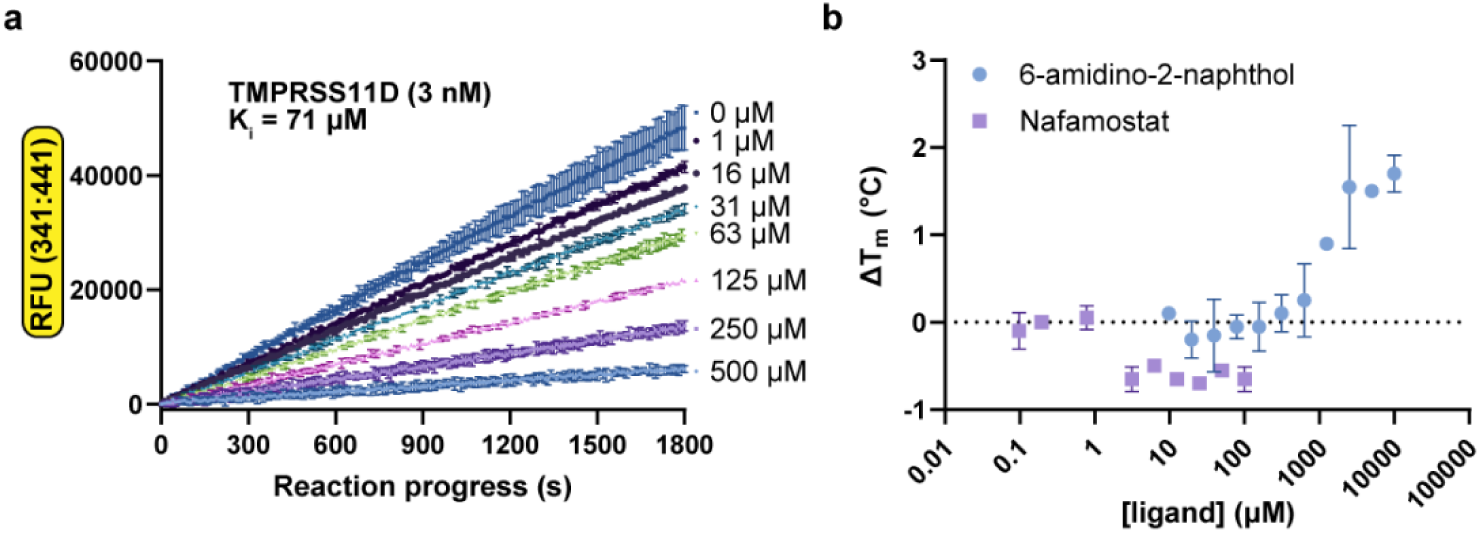
6-amidino-2-naphthol is a noncovalent TMPRSS11D inhibitor that does not depend on the TMPRSS11D catalytic Ser368 residue for binding. **a,** Peptidase activity progress curves of dasTMPRSS11D (3 nM) with 6-amidino-2-naphthol at the indicated inhibitor concentrations added simultaneously with Boc-QAR-AMC substrate (100 µM final). The 6-amidino-2-naphthol apparent inhibition constant (K_i_) was determined through curve-fitting in DynaFit 4.0 (Methods). **b**, Melting temperature shifts (Δ*T*_*m*_s) of dasTMPRSS11D S368A protein in the presence of the indicated concentrations of nafamostat (violet datapoints) or 6-amidino-2-naphthol (teal datapoints) ligands. Each assay contained 2 µg dasTMPRSS11D S368A, 5X SYPRO orange dye, and 50 mM Tris pH 8.0 with 200 mM NaCl. Assays were performed in technical quadruplet (*n=*4) and data are shown as mean +/− s.d.

**Extended Data Figure 4.**
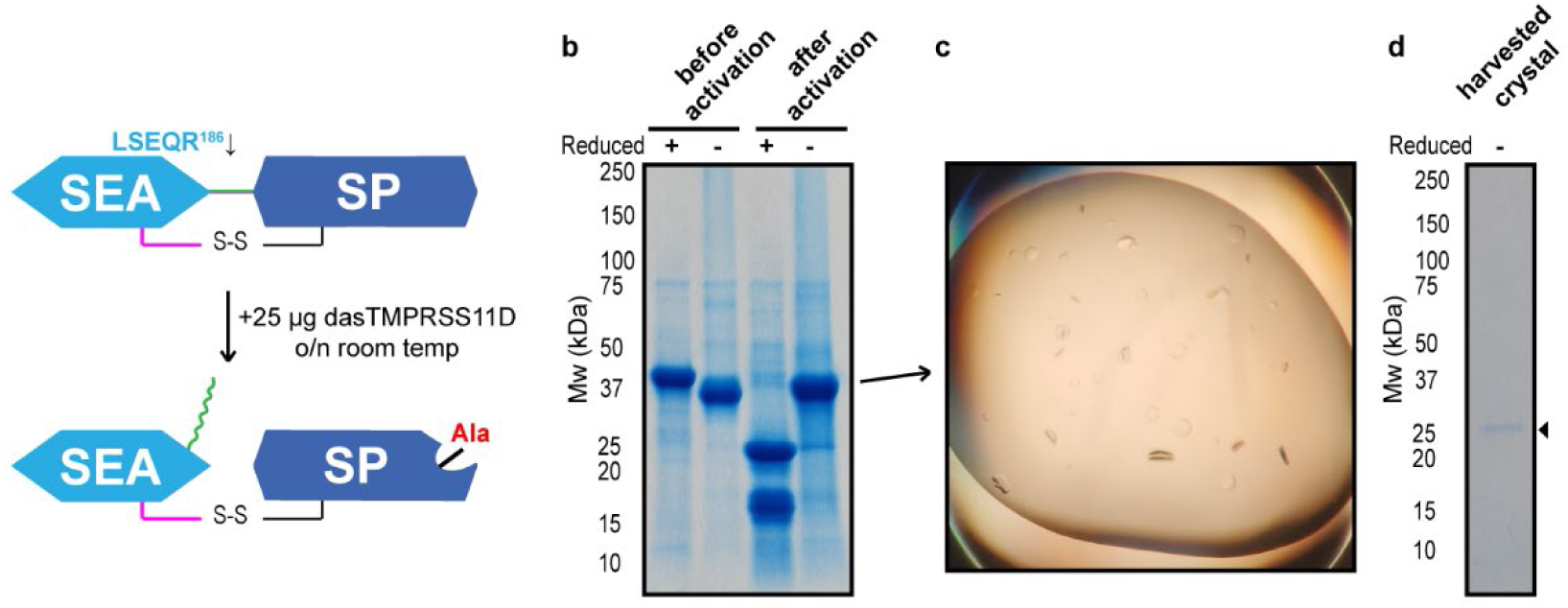
The eTMPRSS11D S368A protein crystal is comprised of a protein lacking the SEA domain. **a**, Schematic of a soluble TMPRSS11D Ser368Ala protein construct before and after treatment with a trace amount of active dasTMPRSS11D protease. **b,** Coomassie-stained SDS-PAGE gel of the eTMPRSS11D S368A protein before and after overnight treatment with 25 µg active dasTMPRSS11D protease. **c**, Protein crystals formed for eTMPRSS11D S368A that provided a high resolution TMPRSS11D crystal structure (PDB 9DPF). **d**, Coomassie-stained SDS-PAGE gel of the eTMPRSS11D S368A protein crystals amenable to structure determine. Crystals were harvested by a loop, washed extensively in fresh precipitant buffer, then prepared for SDS-PAGE through addition of 4X Laemelli buffer.

**Extended Data Figure 5.**
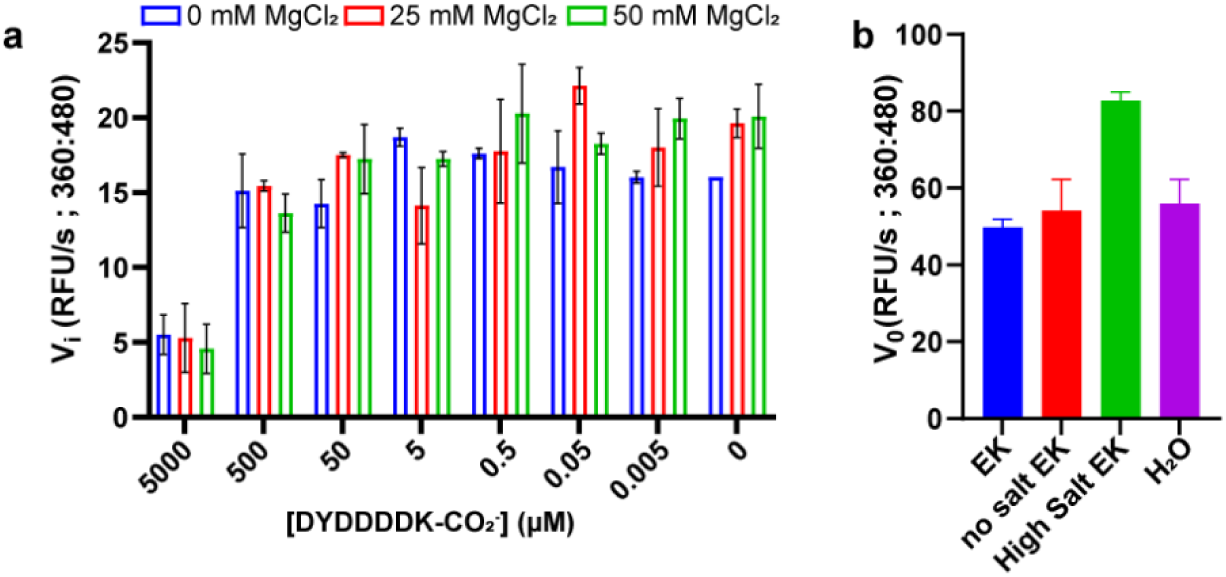
A synthesized DYDDDDK-CO_2_ ^−^ peptide acts as a weak competitive inhibitor of dasTMPRSS11D activity. **a**, Peptidase activity assays in the presence of the indicated concentrations of DYDDDDK-CO ^−^ peptide and MgCl_2_. Approximately 3 nM dasTMPRSS11D protease was used in each assay and protease activity measured using 20 µM Boc-QAR-AMC peptide substrate. Assays were performed in Enterokinase (EK) activity buffer (25 mM Tris pH 8.0, 75 mM NaCl, 2 mM CaCl_2_). Substrate turnover was monitored by fluorescence (360nm:480nm excitation:emission). Initial reaction velocities (V_i_) were measured across the first 120s following substrate addition. **b**, The same peptidase activity assays from (**a**) but contained 0 mM NaCl (no salt EK buffer), 500 mM NaCl (High Salt EK buffer), or were performed in distilled water (H_2_O).

**Extended Data Figure 6.**
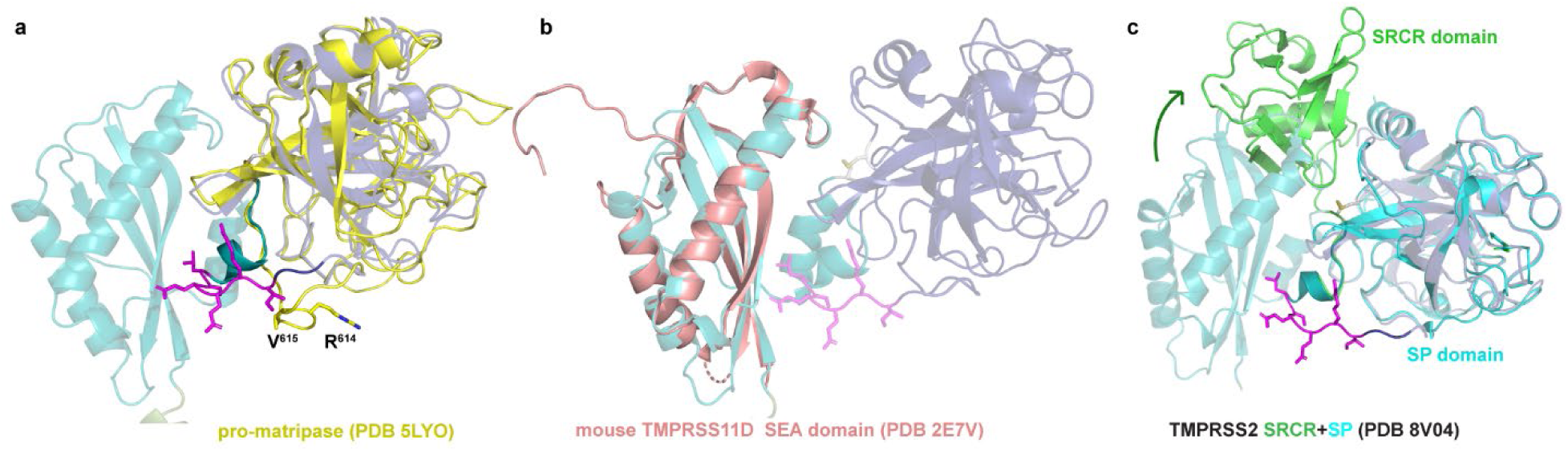
The AlphaFold2 TMPRSS11D structure accurately represents the full-length, zymogen form of the protease. **a**, Structural superposition of the serine protease domain of the AlphaFold2 TMPRSS11D structure (dark purple; AF-O60235-F1) with the pro-matriptase serine protease domain (yellow; PDB 5LYO). The zymogen activation motif of TMPRSS11D is indicated with magenta sticks and the zymogen activation motif of pro-matriptase indicated with yellow sticks. The TMPRSS11D SEA domain is shown as a teal cartoon. **b**, superposition of mouse TMPRSS11D SEA domain (salmon cartoon; PDB 2E7V) upon the SEA domain of the AlphaFold2 TMPRSS11D structure. **c**, Superposition of the TMPRSS2 serine protease domain (teal; PDB 8V04) upon the serine protease domain of the AlphaFold TMPRSS11D serine protease domain (purple; AF-O60235-F1). The SRCR domain of TMPRSS2 is shown as a green cartoon. The relative positioning of the TMPRSS2 SRCR domain to the TMPRSS11D SEA domain is indicated with a green arrow.

**Extended Data Figure 7.**
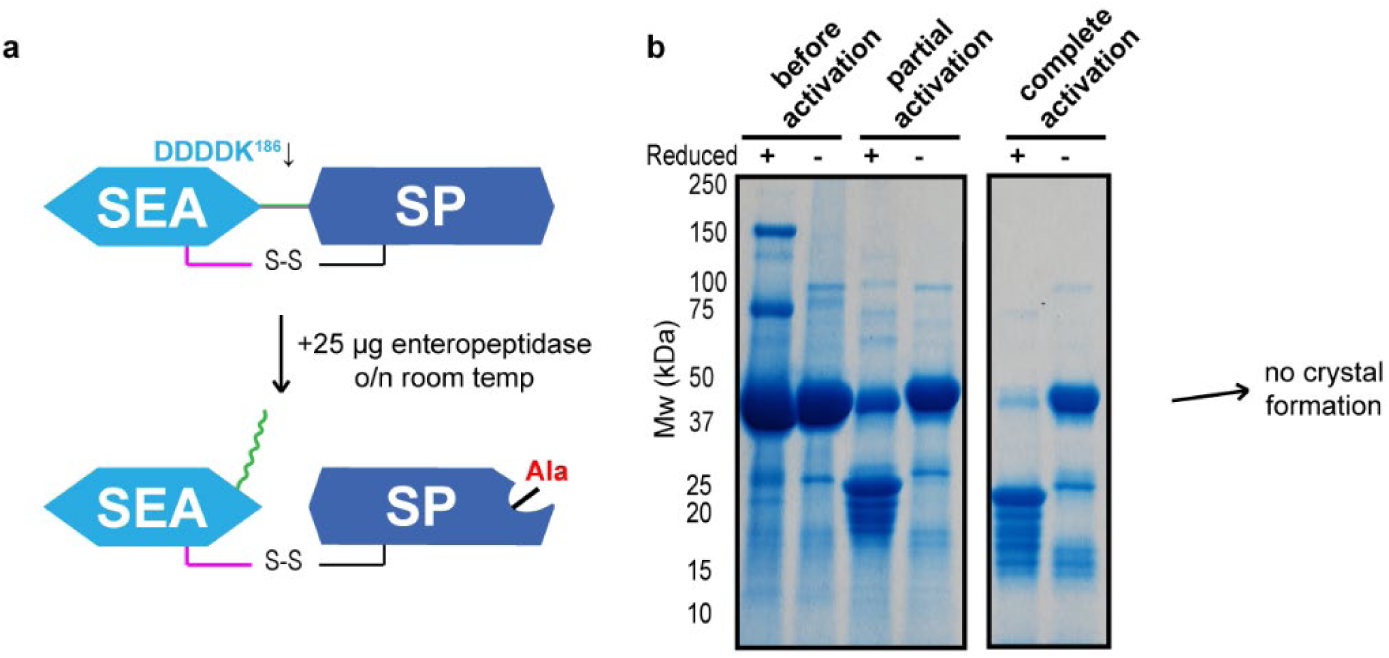
The dasTMPRSS11D S368A protein is amenable to overexpression, purification, and zymogen activation but resists crystallization. **a**, Schematic of a soluble dasTMPRSS11D Ser368Ala protein construct before and after treatment with a trace amount of active human enteropeptidase. **b,** Coomassie-stained SDS-PAGE gel of the dasTMPRSS11D S368A protein before enteropeptidase treatment, 6hrs after enteropeptidase addition (partial activation) and 18 hrs after enteropeptidase addition (complete activation). Approximately 25 µg active human enteropeptidase was used for dasTMPRSS11D S368A protein activation.

