## Supplementary Figure 1 for "Structural basis of TMPRSS11D specificity and autocleavage activation"

### SUPPLEMENTARY INFORMATION

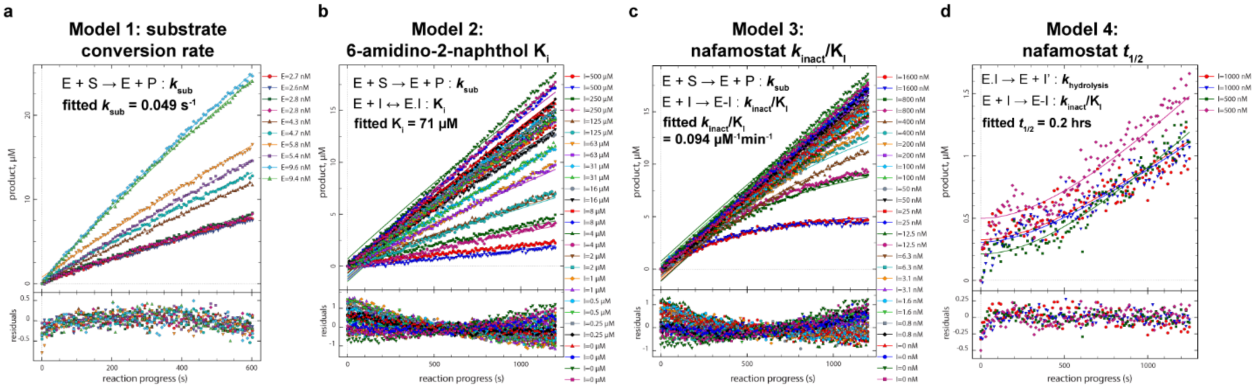


**Supplementary Figure 1. TMPRSS11D enzyme kinetic models to determine kinetic inhibition parameters for 6-amidino-2-naphthol and nafamostat. a**, TMPRSS11D Boc-QAR-AMC substrate conversion rate, $k_{sub}$. Reaction progress curves generated with the indicated concentrations of dasTMPRSS11D enzyme and 100 μM Boc-QAR-AMC substrate and monitored for AMC product formation. Progress curves were fitted across all enzyme concentrations to determine the $k_{sub}$ parameter. Datapoints and curve fits are shown, with data residuals (curve fit value – experimental data) plotted below. **b**, TMPRSS11D inhibition constant, $K_i$, with 6-amidino-2-naphthol. Progress curves were generated using 3 nM dasTMPRSS11D, 100 μM Boc-QAR-AMC substrate, and the indicated concentrations of 6-amidino-2-naphthol. Inhibitor and substrate were simultaneously transferred to wells containing dasTMPRSS11D and fluorescence was monitored immediately. **c**, TMPRS11D inactivation potency parameter, $k_{inact}/K_I$, with nafamostat. Data was generated using 3 nM dasTMPRSS11D and 100 μM Boc-QAR-AMC substrate within the same kinetic assay as (**b**). **d**, TMPRS11D nafamostat inhibition half-life parameter, $t_{1/2}$. Data was generated using 3 nM dasTMPRSS11D and 100 μM Boc-QAR-AMC substrate. Nafamostat (at the indicated concentrations) was incubated with dasTMPRSS11D for 3 minutes prior to substrate transfer. Data was curve-fitted to determine the hydrolysis rate of the dasTMPRSS11D:nafamostat acyl-enzyme complex, $k_{hydrolysis}$. The curve-fitted $k_{hydrolysis}$ parameter was converted to $t_{1/2}$ using $t_{1/2}$=0.693/$k_{hydrolysis}$.
